# FrogCap: A modular sequence capture probe set for phylogenomics and population genetics for all frogs, assessed across multiple phylogenetic scales

**DOI:** 10.1101/825307

**Authors:** Carl R. Hutter, Kerry A. Cobb, Daniel M. Portik, Scott L. Travers, Perry L. Wood, Rafe M. Brown

**Affiliations:** Biodiversity Institute and Department of Ecology and Evolutionary Biology, University of Kansas, Lawrence, KS 66045, USA; Museum of Natural Sciences and Department of Biological Sciences, Louisiana State University, Baton Rouge, LA 70803, USA; Department of Biological Sciences & Museum of Natural History, Auburn University, Auburn, Alabama 36849, USA; California Academy of Sciences, San Francisco, CA 94118, USA

**Keywords:** amphibians, Anura, exon capture, genomics, target capture, UCEs

## Abstract

Despite the increasing use of high-throughput sequencing in phylogenetics, many phylogenetic relationships remain difficult to resolve because of conflict between gene trees and species trees. Selection of different types of markers (i.e. protein-coding exons, non-coding introns, ultra-conserved elements) is becoming important to alleviate these phylogenomic challenges. For evolutionary studies in frogs, we introduce the new publicly available FrogCap suite of genomic resources, which is a large and flexible collection of probes corresponding to ∼15,000 markers that unifies previous frog sequencing work. FrogCap is designed to be modular, such that subsets of markers can be selected based on the phylogenetic scale of the intended study. FrogCap uses a variety of molecular marker types that include newly obtained exons and introns, previously sequenced UCEs, and Sanger-sequencing markers, which span a range of alignment lengths (100–12,000 base pairs). We tested three probe sets from FrogCap using 105 samples across five phylogenetic scales, comparing probes designed using a consensus- or genome-based approach. We also tested the effects of using different bait kit sizes on depth of coverage and missing data. We found that larger bait kits did not result in lowered depth of coverage or increased missing data. We also found that sensitivity, specificity, and missing data are not related to genetic distance in the consensus-based probe design, suggesting that this approach has greater success and overcomes a major hurdle in probe design. We observed sequence capture success (in terms of missing data, quantity of sequence data, recovered marker length, and number of informative sites) and compared them at all phylogenetic scales. The incorporation of different molecular marker types allowed recovery of the variation required for resolving difficult phylogenetic relationships and for performing population genetic studies. Altogether, FrogCap is a valuable and adaptable resource for performing high-throughput sequencing projects across variable timescales.

## Introduction

The widespread adoption of high-throughput DNA sequencing technologies to address questions in evolutionary biology has revealed new challenges for analyzing these expansive genomic datasets (Shendure & Ji 2008; Kircher & Kelso 2010; Jones & Good 2016). Despite the increased ability to collect large phylogenomic datasets, many phylogenetic relationships remain ambiguous because of both biological and non-biological processes that can lead to gene trees that are discordant with the species tree (Knowles 2009; Liu et al. 2009; Hobolth et al. 2011; Blom et al. 2017; Hahn & Nakhleh 2016; Reddy et al. 2017; Knowles et al. 2018; Richards et al. 2018). Further exacerbating this challenge, data from different genomic regions can demonstrate conflicting phylogenomic results (Liu et al. 2009; Lemmon & Lemmon 2013; Chen et al. 2017; Reddy et al. 2017). Despite the decreasing costs of obtaining sequence data, selecting the best sequencing methods and molecular markers for particular projects remains challenging.

Although genome sequencing for non-model organisms remains costly, high-throughput sequencing of targeted genomic regions has resulted in numerous approaches for creating expansive phylogenomic and population genetic datasets at reduced financial costs (Kircher & Kelso 2010; Glenn 2011; Rohland & Reich 2012). These approaches aim to affordably sequence reduced portions of the genome and also allow more individuals and/or species to be sampled simultaneously (Rohland & Reich 2012). An important consideration for systematics and population genetics is balancing the trade-off between obtaining orthologous markers across moderate to deep timescales, while obtaining markers that are variable enough to resolve ambiguous phylogenetic relationships and provide high-quality variants for population genetics (Sulonen et al. 2011; Jones & Good 2016).

There are now many approaches for sequencing subsets of an organism’s genome, and the choice of method largely depends on the phylogenetic breadth required and the number of samples to include. The most common methods for obtaining subsets of genome-wide sequence data include: (1) restriction-associated digestion methods (RADseq) that target markers adjacent to restriction enzyme sites (Miller et al. 2007); (2) targeted sequence capture (HybSeq) methods that target genomic regions through hybridization-based sequence capture (Hancock-Hanser et al. 2013; Jones & Good 2016), including exon capture; and (3) transcriptome sequencing (RNASeq) that target the expressed exome of a sampled tissue type (Wang et al. 2009). Exome capture is another approach similar to HybSeq except it is highly specialized for a reduced taxonomic scale, because probes are designed from transcriptomes/genomes within a particular focal group (Choi et al. 2009; Sulonen et al. 2011). Each of these three methods have been used to address a variety of phylogenetic and population genetic questions and have method-specific advantages and disadvantages (reviews in Lemmon & Lemmon 2013 and McCormack et al. 2013), including tradeoffs regarding costs, phylogenetic scale, and availability of initial genomic resources necessary for study design and development.

Phylogenomic data obtained from each of these methods has the potential to resolve difficult phylogenetic relationships, yet many relationships remain ambiguous. Numerous studies suggest that natural processes such as incomplete lineage sorting, horizontal gene transfer, gene loss or duplication, and hybridization can lead to gene trees that are discordant with the species tree (Maddison 1997; Liu et al. 2009; Knowles 2009; Knowles et al. 2018). Additionally, natural selection of protein coding and other selected regions can be problematic for phylogenetic inference if phylogenetic signal is reduced as a result of selection (Castoe et al. 2009; Edwards 2009; Liu et al. 2009; Hobolth et al. 2011). Discordant relationships can also be caused by non-biological mechanisms, such as gene tree estimation error resulting from model inadequacy, short alignment lengths, low levels of phylogenetic informativeness, anomaly zones, or errors in sequence assembly or alignment (Xi et al. 2015; Hahn & Nakhleh 2016; Blom et al. 2017; Reddy et al. 2017; Richards et al. 2018). As a result, selecting the best method and types of molecular markers to address these issues remains challenging.

Selection of different types of molecular markers (e.g. protein-coding exons, non-coding regions, ultra-conserved elements [UCEs], or single nucleotide polymorphisms [SNPS]) is an increasingly important issue when dealing with the challenges of analyzing large phylogenomic datasets. Each phylogenomic marker type may be useful at various timescales and may present different biases for analyses (Lemmon & Lemmon 2013; McCormack et al. 2013; Hosner et al. 2016). For example, there is debate over whether protein-coding or non-coding markers are more appropriate for phylogenetic analyses, because selection on protein-coding exons may mask homology and bias phylogenetic signal (Liu et al. 2009; Chen et al. 2017; Reddy et al. 2017). One well-studied case in birds exemplifies this debate, where conflicting relationships inferred from different phylogenomic studies have produced non-trivial differences in phylogeny from the usage of different marker types (McCormack et al. 2013; Prum et al. 2016; Reddy et al. 2017). In addition, other marker features such as locus length (Edwards et al. 2016; Springer & Gatesy 2016) or character type (e.g. SNPs: Leaché & Oaks 2017; insertions or deletions: Simmons & Ochoterena 2000; transposable elements: Han et al. 2011) have been explored to find optimal data types for phylogenomic and population genetic analyses.

These debates have made it clear no consensus exists regarding a perfect, universal molecular marker type, capable of simultaneously alleviating all of these problems. As such, choice of marker type remains a substantial challenge, which presents numerous obstacles for both existing and future phylogenomic datasets. One solution is to use a diverse assortment of markers sampled from across the genome with different properties (Dool et al. 2016; Chen et al. 2017). The inclusion of a diversity of marker types allows downstream filtering of markers by desirable properties, such as length or informativeness, which can improve phylogenetic estimates (Mirarab et al. 2014; Springer & Gatesy 2016; Chakrabarty et al. 2017; Streicher et al. 2018; Karin et al. 2019). When considering these factors, using a HybSeq approach that targets a variety of marker types may represent an optimal solution; however, substantial genomic resources are required for the initial probe design for the target markers.

Targeted capture using HybSeq is widely used by the phylogenetics community and has been dominated by targeting and sequencing conserved elements through two main approaches: anchored hybrid enrichment using conserved exons (AHE, Lemmon et al. 2012) and ultra-conserved elements (UCEs, Faircloth et al. 2012). These approaches identify regions of the genome that are conserved across distantly related taxa and use probes designed from these conserved regions, which are used in hybridization-based target capture (Gnirke et al. 2009). The two methods differ in that AHE targets ∼400 medium length exons (>500bp) that are moderately divergent but sufficiently conserved to capture regions across hundreds of millions of years (Lemmon et al. 2012). In contrast, the UCE approach targets ultra-conserved regions (∼120 bp long) with the goal of obtaining highly variable flanking regions adjacent to the conserved regions (Bejerano et al. 2004; Faircloth et al. 2012). Both types of markers have been criticized, where AHE markers have the potential to bias phylogenetic results due to natural selection (AHE targets only exons, Castoe et al. 2009; Bragg et al. 2016; Singhal et al. 2017). In addition, UCE markers have been criticized for their unknown function (Alexander et al. 2010), low variation in many UCEs (McCormack et al. 2012), and difficulties related to aligning highly variable flanking regions among distantly related taxa (Singhal et al. 2017; Streicher et al. 2018). Both AHE and UCEs are now widely used in inferring phylogenies across broad and shallow phylogenetic scales (Crawford et al. 2012; Smith et al. 2014; Brandley et al. 2015; Prum et al. 2015; Streicher et al. 2018). For population genetics, UCEs have been used with some success (Hosner et al. 2016; Zarza et al. 2016; Andermann et al. 2019). In contrast, AHE produces a smaller number of markers (less than 400 exons) and generally results in fewer independent variants as required by many types of analyses in comparison to these other sets of markers (Lanier & Knowles 2012; Hedge & Wilson 2014; Springer & Gatesy 2018). Despite the widespread usage of phylogenomic data, no single probe set composed of multiple marker types exists that can be employed across a wide range of organisms.

Amphibians represent one of the most diverse terrestrial vertebrate groups, and much of this diversity is concentrated within a single lineage – frogs (Anura). Frogs have been diversifying for over 200 million years and now include over 7000 species (88% of amphibians; AmphibiaWeb 2019). Because of this deep time scale and numerous rapid radiations across the diversity of frog clades, this exceptional diversity has presented a major challenge for research into amphibian evolution and the performance of custom targeted sequence capture remains largely unexplored (but see Hedtke et al. 2013; Portik et al. 2016; Salamanders: McCartney-Melstad 2016). Several studies have used UCEs (Alexander et al. 2017; Pie et al. 2018; Streicher et al. 2018; Zarza et al. 2018; Guillory et al. 2019) and others have used AHE markers (Peloso et al. 2016; Heinicke et al. 2018; Yuan et al. 2018), whereas two additional studies created customized probe sets for an African frog clade (Afrobatrachia) and the Asian genus *Limnonectes* (Portik et al. 2016; Reilly et al. 2019). Although UCEs have been used in frogs, they are not ideal because they were designed for amniotes, a group which does not contain frogs; therefore, about half the target UCEs are typically captured (∼2500/5600 UCEs; Streicher et al. 2018; Guillory et al. 2019). The AHE probe set is advantageous because it produces long exons that can be more predictably modelled. However, longer exons require more probes for suitable capture efficiency, and this reduces the number of exons that can be included (Bragg et al. 2016; Singhal et al. 2017). In summary, despite the widespread use of high-throughput sequencing in non-model organisms and the increasing number of studies in frogs, a probe set incorporating different molecular marker types for the substantial diversity across the global frog radiation has not yet been made available.

We introduce FrogCap, a publicly available collection of molecular markers for all frogs that can be adapted into sequence capture probe sets that includes markers from a variety of data types (exons, introns, UCEs, markers sequenced previously using traditional/Sanger methods, independent markers to extract variants). We provide a modular, large, and flexible collection of tested markers and probes corresponding to ∼15,000 markers, which unifies previous sequencing via the inclusion of “legacy” Sanger sequencing markers traditionally used in phylogenetic studies (Figure 1; Frost et al. 2006; Pyron & Wiens 2011; Feng et al. 2017) and UCEs that have been successfully captured in anurans (Alexander et al. 2016; Streicher et al. 2018). The FrogCap marker set is designed to be modular, such that subsets of the markers can be selected based on the probe set size, type of research question, and the taxonomic scale being addressed. We describe a novel approach for selecting sets of orthologous markers that functions well across divergent taxa within and across the two main superfamilies of frogs (Hyloidea + Archaeobatrachia and Ranoidea), by creating two complementary probe sets (referred to as the Hyloidea and Ranoidea probe sets), which allow markers to be combined across these two superfamilies (Feng et al. 2017). We also test the modularity of FrogCap by creating a third probe set, which is a reduced version of the Ranoidea set that uses half the number of probes (termed Reduced-Ranoidea hereafter), to evaluate whether the reduction in probes leads to increased depth of coverage and higher quality variant data. Finally, we provide a new bioinformatics pipeline to analyze sequence capture data, which begins with raw demultiplexed sequence data and produces cleaned alignments and variant datasets.

**Figure 1.**
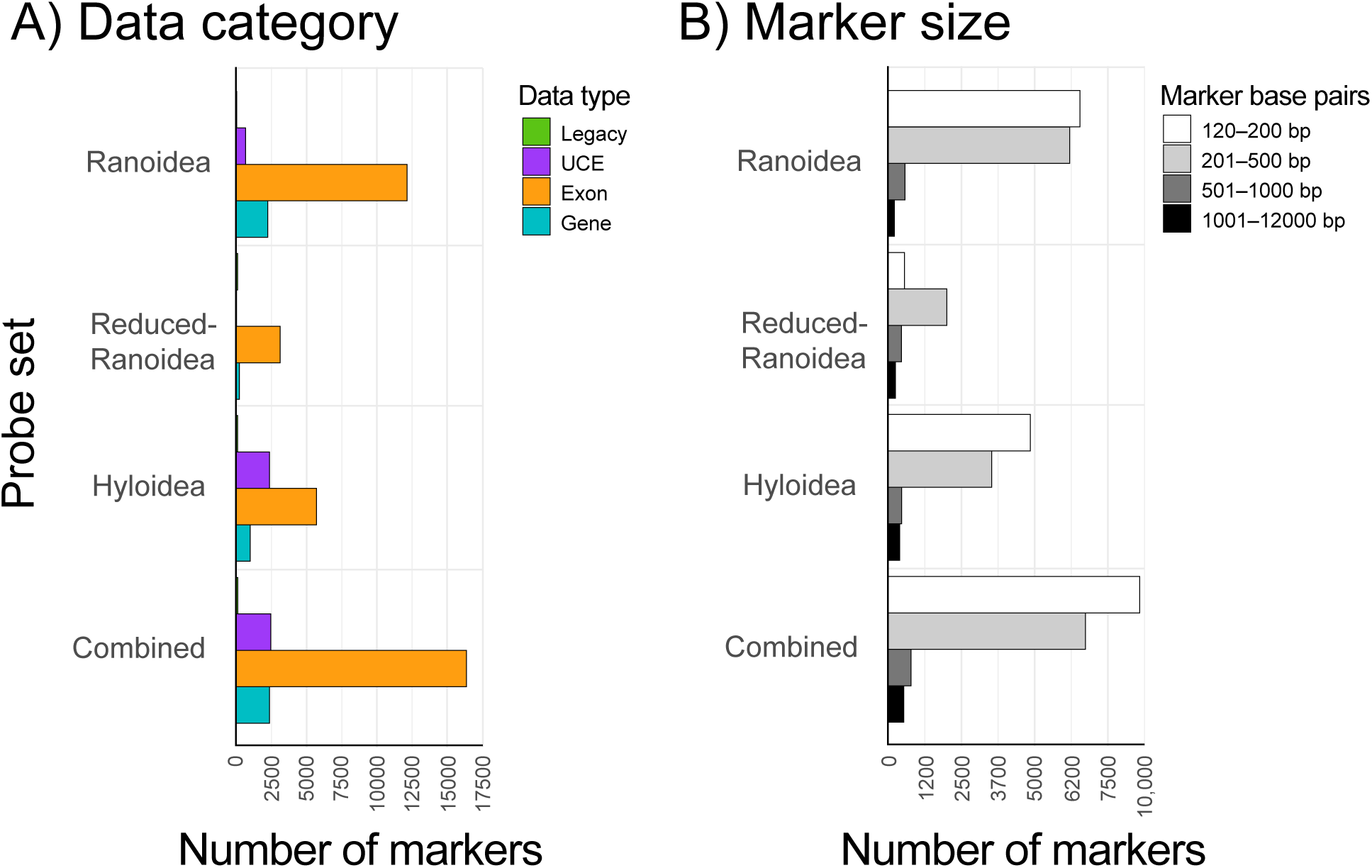
The modularity of the FrogCap probe set permits the selection of different types of markers. The data category in A) shows the quantity of different marker types (Legacy, UCE, Exon, Gene) used in the design of each probe set (40K bait set used for Ranoidea and Hyloidea; 20K bait set used for Reduced-Ranoidea). The marker size distribution in B) shows the general size classes of markers used for each of the probe sets. The “Combined” probe set refers to the number of unique markers across all probe sets to represent the total available markers for FrogCap. Different configurations can be created based on bait kit size, modules, marker size or phylogenetic group.

Using new sequence data, we test the utility of these three probe sets using 105 samples and evaluate the sensitivity and specificity of the probe design, the number of markers captured, and depth of coverage across the different probe sets. We use the resulting alignments from each phylogenetic scale to assess the number of markers and alignment lengths recovered, the proportion of informative sites, and levels of missing data. Finally, we examined the effects of phylogenetic relatedness on our capture success, assessing how genetic distance relates to missing data, sensitivity, and specificity of the sequence capture. Lastly, we test whether using fewer probes in the Reduced-Ranoidea set leads to greater capture success, increased depth of coverage, and higher quality variant data.

## Materials and Methods

### FrogCap website

We have developed a website (https://www.frogcap.com) that can be used to configure and select different probe sets using the FrogCap marker database. The webpage and Github page (https://github.com/chutter/FrogCap-Sequence-Capture) provide the pre-configured probe sets developed and tested in this study. In addition, the configuration website will allow the selection of different clades of interest for differently sized bait sets, as well as well selection of different marker types (exons, introns, UCEs, legacy) and different size classes (under development).

### Sequence Capture Probe Design

#### Transcriptome assembly

Thirty anuran transcriptomes were obtained from previously published studies (Supplementary Table S1). Raw reads were downloaded from the NCBI sequence read archive using the SRA Toolkit for 26 transcriptomes (Leinonen et al. 2010). Adapter sequences were trimmed with PEAT (Li et al. 2015) and reads were quality trimmed and filtered with AfterQC (Chen et al. 2017). Trimmed and filtered reads were assembled with Bridger under default settings (Chang et al. 2015). We also included four additional transcriptomes of Portik et al. (2016), using sequencing and assembly methods described therein. After assembly, isoforms, alternatively spliced sequences, and other redundancies were removed using CD-HIT-EST (Li & Godzik 2006) at a similarity threshold of 95%, keeping the longest open reading frame. These reduced-redundancy transcriptomes were used for subsequent probe design.

**Table 1.** Marker contents targeted for three probe sets designed in this study. The Reduced probe set uses the same markers and probes from the Ranoidea set but was designed with half the number of baits for a specific taxonomic group. Ranoidea V2 was not explicitly used in this paper; however, it represents a revision of Ranoidea V1 where failed markers were discarded in favor of additional UCEs and Legacy markers.

|  | **Hyoidea** | **Ranoidea V1** | **Ranoidea V2** | **Reduced** |
| --- | --- | --- | --- | --- |
| *N* probes | 40,040 | 40,040 | 40,040 | 20,020 |
| *N* base pairs targeted | 2,929,956 | 3,454,114 | 3,313,548 | 1,519,233 |
| *N* markers targeted | 9,229 | 13,517 | 12,909 | 3,247 |
| Hyoidea overlap | - | 6,130 | 6,931 | 1,652 |
| Ranoidea V1 overlap | 6,130 | - | 10,476 | 3,136 |
| Ranoidea V2 overlap | 6,931 | 10,476 | - | 2,953 |
| Reduced overlap | 1,652 | 3,136 | 2,953 | - |
| *N* exons | 6,977 | 12,834 | 10,743 | 3,161 |
| *N* UCEs | 2,166 | 651 | 2,080 | 0 |
| *N* Legacy | 86 | 32 | 86 | 86 |
| Mean marker length | 317.2 | 255.5 | 256.7 | 467.8 |
| Maximum marker length | 11,549 | 11,429 | 11,429 | 11,429 |
| Markers < 500 bp | 8,382 | 12,733 | 12,222 | 2,554 |
| Markers 500-1000 bp | 2,562 | 616 | 441 | 447 |
| Markers > 1000 bp | 390 | 168 | 246 | 246 |

#### Ranoidea genome-based probe design

To target new exons for sequence capture, we designed a probe set by locating orthologous, protein-coding exons that were well represented across frogs of the superfamily Ranoidea. We used the *Nanorana parkeri* genome (Sun et al. 2015) and that study’s original genomic annotations, to determine the genomic coordinates of predicted exons in the genome (genome and annotations available at GigaScience: dx.doi.org/10.5524/100132). We conducted all data processing and other analyses in R (RDCT 2018), using customized scripts with the following R packages: GENOMICRANGES (Lawrence et al. 2013), SEQINR (Charif & Lobry 2008), and APE (v5.0; Paradis & Schliep 2019).

First, we used the program BLAT (Kent 2002) to match each assembled frog transcriptome to the *Nanorana* genome using a 75% similarity threshold. Second, we combined the match dataset for each species into a single dataset, such that orthologous matches could be conveniently clustered together (this resulted in 2.7 million match records). Third, we filtered out matches that were less than 50 bp and matched to multiple locations in the genome, to remove potential paralogs (removed ∼2 million of the aggregated matches; 749,821 matches remained). Fourth, matches were clustered together when they overlapped within a predicted exon from the *Nanorana* genome (removed ∼500,000 matches; 228,960 matches remained). After these initial filtration steps of the matches, a final candidate marker set of 94,293 exons remained.

The sequence data for each candidate exon match above were collected from each transcriptome and aligned together with the *Nanorana* genome exon sequence using MAFFT v7.312 (auto and default parameters; Katoh & Stanley 2013). To assist in exon selection, each multiple species alignment was assessed and filtered through the following criteria: (1) kept alignments with >5% parsimony informative sites to ensure some variability was present; (2) kept alignments with GC content between 30–50%, because such sequences are efficient for sequence capture (Gnirke et al. 2009); (3) exons <100BP and <5000bp were removed to include more exons from across the genome; (4) exons were kept only if they were found in at least 50% of the species transcriptomes; and (5) potential paralogs were removed by using BLAT to detect multiple hits across both *Nanorana* and *Xenopus tropicalis* (Hellsten et al. 2010) genomes. After exon assessment and removal filtration steps, 18,502 exons remained in the final candidate exon dataset, with a mean length of 152 bp and a mean pairwise divergence of 13.6 percent.

The *Nanorana* genome sequence was extracted for each candidate exon, and the genome sequence was used to design a MYbaits-2 (40,040 baits) custom bait library (MYcroarray, now Arbor Biosciences), using 120mer baits to best capture sequences with greater than 5% divergence from the probes. For the UCE markers, we used a subset of the 120mer UCE Amniote probes from Faircloth et al. 2012 (https://www.ultraconserved.org). To design remaining probes from the *Nanorana* genome sequences, the finalized set of markers were separated into probe sequences following a 2x tiling scheme, starting 20bp behind the start codon of the exon and tiling 120bp probes every 60bp until 20bp past stop codons (total: 61,939 baits). Individual probes were filtered using these criteria: (1) excluded probes that matched to multiple locations in both *Nanorana* and *Xenopus* genomes with BLAT with a 70% similarity; (2) kept probes with a GC content between 30–50%; (3) kept if there were no repetitive sequences based on the *Nanorana* RepeatMasker annotations (Sun et al. 2015); and (4) kept if there were no matches with BLAT to other probes (using a 70% similarity criterion). After filtration 51,344 baits remained; markers and their baits were randomly selected to drop 11,304 baits from the dataset to fit into the 40,040 bait limit. This resulted in a final set of 12,834 exons, which we combined with 32 legacy markers and 651 UCEs (selection described below). The final Ranoidea V1 probe set included 13,517 markers covering a total of 3,454,114 bp (Table 1; Figure 1). After testing the Ranoidea V1 probe set in this study, we make available a revised Ranoidea V2 probe set that was created that excluded unsuccessful markers and included the 64 additional legacy Sanger markers from Feng et al. 2017 (including the 32 above; see below) and longer exons (see Reduced-Ranoidea design; Table 1). All testing results in this study are derived from the Ranoidea V1 probe set (referred to as Ranoidea onward).

#### Hyloidea transcriptome-based probe design

To design probes from orthologous markers for frogs from the Hyloidea superfamily, there was no available genome for this group, so a different approach was required. We began with the Ranoidea exons designed above and attempted to match exons to six hyloid transcriptomes using BLAT (Supplementary Table S1). We clustered together matches, created multiple sequence alignments, and filtered as described above for the genome-based probe design. After these steps, the candidate dataset included 6,130 markers. For these matching exons, we generated consensus sequences using the transcriptomes, and ambiguous sites were replaced with a base that would maintain an optimal GC content.

To add additional exons to this dataset, we used VSEARCH (Rognes et al. 2016) to cluster together transcripts from the Hyloidea transcriptomes via the identification of orthologous clusters (18,949 cluster were found). We obtained the consensus sequence for each cluster and matched it back to each transcriptome (resulting in 101,863 matches). Next, we generated 18,949 sequence alignments for each cluster and filtered the alignments as described for the Ranoidea V1 set. We removed any alignments with two or less samples, which resulted in 16,511 transcript alignments with 3–6 samples per alignment. We randomly selected 926 transcripts (mean length: 677.8 bp; range: 153–11,188 bp) to compliment the 6,130 shared redesigned Ranoidea markers (described above), and the 2,166 UCEs and 86 Feng et al. (2017) legacy Sanger markers described below. We created the final set of markers to be used in probe design by generating consensus sequences from each filtered transcript alignment. Finally, we designed and filtered probes as described above for the Ranoidea V1 set using the MyBaits-2 kit of 40,040 baits. The final Hyloidea probe set targeted 9,229 markers covering a total of 2,929,956 bp (Table 1; Figure 1).

#### Reduced-Ranoidea marker selection

To test the modularity and customization potential of FrogCap, we selected a reduced set of markers from the 40,040-bait kit, for use in our smaller, 20,020-bait kit. For this probe set, we also tested additional markers that could be incorporated into future customized kits. We included the 86 Feng et al. (2017) markers described in the next section, which were included in the Hyloidea and Ranoidea V2 (but not the Ranoidea V1) set. We also included 47 ultra-long exons (>5000bp) previously not included in the Ranoidea V1 set. These ultra-long exons also have been included in the Ranoidea V2 set (Table 1).

The Reduced-Ranoidea set was designed after the 40K Ranoidea V1 set had been tested, which enabled the selection of markers that were captured successfully across the entire superfamily. To select markers that had already been tested, we began with alignments from the 24 sample Ranoidea set evaluated below, and reduced the probe set to markers successfully captured across 75% or greater of the samples, which was 10,274 markers. Next, we filtered the markers as follows: (1) UCEs were excluded; (2) the largest exon within 100,000 bp of another exon on the *Nanorana* genome was kept reducing potential genetic linkage due to recombination; and (3) all exons greater than 500 bp were included. This final candidate set included ∼4,312 exons. To accommodate the probe limit, exons were randomly deleted until the 20,020 bait limit was reached, resulting in 3,161 exons retained. After combining the 86 legacy markers, our final Reduced-Ranoidea marker set included 3,247 markers targeting 1,519,233 bp of data (Table 1; Figure 1).

#### Previously published legacy markers

To maintain compatibility with previously published datasets, we assessed 40 commonly used nuclear markers for phylogenetic studies in frogs (e.g. Frost et al. 2006; Pyron & Wiens 2011). The selected legacy markers were mapped to the *Xenopus* and *Nanorana* genomes, ensuring that none were paralogs or had portions matching to multiple genomic regions. We selected 36 markers from this assessment; this set of legacy markers were incorporated in the Ranoidea V1 probe set by extracting these markers from the *Nanorana* genome sequence (Figure 1). To include markers from a more recent study in the Hyloidea and Reduced-Ranoidea probe sets (that were not published prior to the development Ranoidea V1), we assessed the 95 markers from Feng et al. (2017). We removed any duplicates overlapping with the legacy marker set (36 markers) and again mapped all markers to the two reference genomes. Our combining and condensing of these two sets of legacy markers resulted in a final set of 86 legacy markers. Final marker sequences were designed from the consensus sequences across the multiple sequence alignments from Feng et al. (2017) and were then used for our probe design. Although the Feng et al. (2017) markers were not available for the Ranoidea V1 probe set, they are included in the revised Ranoidea V2 probe set (Table 1).

Ultra-conserved elements have been previously used in frogs, with ∼50% success capture rate (e.g. Alexander et al. 2016; Pie et al. 2018; Streicher et al. 2018). For this study, we selected a subset of the orthologous UCEs previously sequenced from *Kaloula* (Microhylidae: Alexander et al. 2016). We selected a subset of UCEs that had greater than 10% parsimony informative sites, for a total of 651 UCEs included in Ranoidea V1 using stock UCE probe sequences. Improving upon this for the Hyloidea set, we included the 2,166 successfully captured UCEs from Streicher et al. (2018), which contains the 651 UCEs selected above. For these UCEs, we redesigned our probe sequences by creating consensus sequences across the multiple sequence alignments for each UCE from Streicher et al. (2018).

#### Marker overlap

An important design component of these complimentary probe sets is that they contain markers that overlap with each other (Table 1). In the Hyloidea and Ranoidea probe sets, markers that overlap were designed specifically for those superfamilies: in Ranoidea, the probes were designed from the *Nanorana* genome sequences while in Hyloidea the overlapping markers were designed from consensus sequences of the transcriptomes from that superfamily. Hyloidea and Ranoidea V1 overlap with 6,140 markers and Ranoidea V2 overlaps with 6,931 markers from Hyloidea, with the increase coming from a shared 2,068 UCEs (Ranoidea V1 contains 651 UCEs).

### Sequencing

#### Taxonomic sampling strategy

To test these new sequence capture probe sets, we aimed to explore their performance across seven different taxon sampling strategies which spanned five phylogenetic scales. We selected 105 samples for this study and we assessed the following phylogenetic scales: (1) Order, we included 48 samples including samples using the Ranoidea and Hyloidea sets, representing Anura; (2) Superfamily, we assess the Ranoidea and Hyloidea superfamilies independently, including 24 samples from each superfamily (the same samples used in the Order assessment; Hyloidea also tests out one Salamander and two Archaeobatrachrian frogs that are not included in the Hyloidea superfamily); (3) Family, we included eight samples from the family Mantellidae, which include a single representative for each genus except two monotypic genera (Glaw & Vences 2006); (4) Genus, we included samples of 24 species from the genus *Cornufer* (Ceratobatrachidae), which includes representatives from each of the recognized subgenera within this genus (Brown et al. 2015); and (5) Species, we include 16 samples from within a single species, *Cornufer vertebralis*, from four different Solomon Islands insular populations. We also evaluated the Reduced-Ranoidea set using 30 Philippine samples from the genus *Occidozyga* (Dicroglossidae; sampled from most major landmasses throughout the archipelago), to represent a dataset comparable to our Genus dataset. See Supplementary Table S2 for detailed sampling information.

**Table 2.** Sequence capture evaluation for each probe set. Our Reduced probe set uses the same markers and probes from the Ranoidea set but was designed with half the number of baits for a specific taxonomic group. “X” depth refers to the number of overlapping bases in a given marker.

|  | **Hyloidea** | **Ranoidea** | **Reduced** |
| --- | --- | --- | --- |
| *N* probes | 40K | 40K | 20K |
| *N* base pairs | 2,929,956 | 3,454,114 | 1,519,233 |
| *N* markers | 9,229 | 13,517 | 3,247 |
| Mean sensitivity (%) | 70.3% | 70.6% | 89.1% |
| Mean specificity (%) | 47.9% | 71.1% | 35.7% |
| Mean marker missing data (%) | 21.9% | 22.7% | 4.5% |
| Median depth of coverage | 19.7X | 22.1X | 26.6X |
| Median exon depth | 40.4X | 43.6X | 48.1X |
| Median intron depth | 13.8X | 13.2X | 15.8X |

#### Library preparation and sequencing

Genomic DNA was extracted from the 105 tissue samples using either a standard phenol-chloroform extraction or through the use of a Promega^TM^ Maxwell bead extraction robot. The resultant DNA was quantified using a Promega Quantus^TM^ fluorometer. Approximately 500 ng total DNA was acquired and set to a volume of 50 ul through dilution (with H20) or concentration (using a vacuum centrifuge) of the extraction when necessary.

The genomic libraries for the samples were prepared by MYcroarray library preparation service. Prior to library preparation, the genomic DNA samples were quantified with fluorescence and up to 4 µg was then taken to sonication with a QSonica Q800R instrument. After sonication and SPRI bead-based size-selection to modal lengths of roughly 300 bp, up to 500 ng of each sheared DNA sample were taken to Illumina Truseq-style sticky-end library preparation. Following adapter ligation and fill-in, each library was amplified for 6 cycles using unique combinations of i7 and i5 indexing primers, and then quantified with fluorescence. For each capture reaction, 125 ng of 8 libraries were pooled, and subsequently enriched for targets using the MYbaits v 3.1 protocol. Enrichment incubation times ranged 18–21 hours. Following enrichment, library pools were amplified for 10 cycles using universal primers and subsequently pooled in equimolar amounts for sequencing. Samples were sequenced on an Illumina HiSeq 3000 with 150 bp paired-end reads with 96 samples sequenced per lane of sequencing.

### Data processing and alignment

#### Data processing pipeline

A bioinformatics pipeline for filtering adapter contamination, assembling contigs, and exporting alignments is scripted in R and available at (https://github.com/chutter/FrogCap-Sequence-Capture). Prior to processing raw reads, Illumina sequence data were de-multiplexed using the Illumina software bcl2fastq. Next, raw reads were cleaned of adapter contamination, low complexity sequences, and other sequencing artifacts using the program FASTP (default settings; Chen et al. 2018). Adapter-cleaned reads were then matched to a database of bacterial (human skin, ultra-pure water contamination, and other common bacteria; Laurence et al. 2014) and other genomes (*C. elegans, Drosophilia*) to ensure that no contamination persisted in our final dataset (see Supplementary Table S3 for GenBank Accession Numbers of reference genomes). We decontaminated the adapter-cleaned reads with the program BBMAP from BBTools (https://jgi.doe.gov/data-and-tools/bbtools/), by matching cleaned reads to each reference contaminant genome (reads removed if they matched >95 percent similarity). After this step, final reads were saved separated as “cleaned-reads,” which were processed for subsequent variant calling (described below).

**Table 3.** Summary of the All-Markers dataset for each phylogenetic scale. The abbreviation PIS refers to parsimony informative sites.

| Clade<br>(Scale) | Total<br>samples | N samples<br>(mean ± sd) | N<br>markers | Total bp | Length | Total PIS | PIS<br>(mean ± sd) |
| --- | --- | --- | --- | --- | --- | --- | --- |
| Anura<br>(Order) | 48 | 35 ± 8 | 5500 | 1,806,209 | 328.4 ±<br>189.5 | 944,195 | 52.5 ± 10.6 |
| Hyloidea<br>(Superfamily) | 24 | 17.1 ± 5.3 | 8712 | 4,678,657 | 537 ±<br>312.4 | 1,432,769 | 31.9 ± 12.8 |
| Ranoidea<br>(Superfamily) | 24 | 17.1 ± 5 | 13099 | 5,421,176 | 413.9 ±<br>202.8 | 2,144,350 | 40.6 ± 11.5 |
| Mantellidae<br>(Family) | 8 | 7 ± 1.1 | 12106 | 8,129,529 | 671.5 ±<br>249.5 | 818,032 | 10.2 ± 4.6 |
| *Cornufer*<br>(Genus) | 24 | 20 ± 4.8 | 12701 | 7,526,490 | 592.6 ±<br>206.1 | 1,032,575 | 13.8 ± 5.6 |
| *Cornufer*<br>*vertebralis*<br>(Species) | 16 | 14.1 ± 2.9 | 12009 | 7,061,983 | 588.1 ±<br>182.4 | 155,675 | 2.2 ± 2.5 |
| *Occidozyga*<br>(Reduced) | 30 | 27 ± 5.3 | 3156 | 2,268,852 | 718.9 ±<br>545.2 | 243,238 | 11.4 ± 4.6 |

Prior to assembly, the “cleaned-reads” were further processed to decrease computational load and increase accuracy. We merged paired-end reads using BBMerge (Bushnell et al. 2017) from BBTools. BBMerge also fills in missing gaps between non-overlapping paired-end reads by assembling missing data from the other paired-end reads using the “Tadpole” program. Next, exact duplicates were removed if both read pairs were duplicated, using “dedupe” from BBTools. Additionally, duplicates from the set of merged paired-end contigs were removed if they were exact duplicates or were contained within another merged contig.

Merged singletons and paired-end reads were assembled *de novo* using the program SPADES v.3.12 (Bankevich et al. 2012), which runs BAYESHAMMER (Nikolenko et al. 2013) error correction on the reads internally. Data were assembled using several different k-mer values (21, 33, 55, 77, 99, 127), in which orthologous contigs resulting from the different k-mer assemblies were merged. We used the DIPSPADES (Sofanova et al. 2015) function to assemble contigs that were polymorphic by generating a consensus sequence from both haplotypes from orthologous regions such that polymorphic sites were resolved randomly.

Consensus haplotype contigs were then matched against reference marker sequences used to design the probes separately for our three probe sets with BLAST (*dc-megablast*). Contigs were discarded if they failed to match ≥ 30% of the reference marker, and contig matches fewer than 50 bp were removed. Contig matches to a given reference marker were discarded if more than one contig matched to the marker and were overlapping. For non-overlapping matches to the same reference marker, we merged these contigs by joining them together (Ns inserted in matching positions). The final set of matching contigs were labelled with the name of each marker, followed by each sample’s unique institutional identifier (i.e., the corresponding museum voucher catalog number, field collector number, etc.), and assembled in a single file to be parsed out separately for multiple sequence alignment in the next step.

#### Alignment and trimming

Next, our final set of matching markers was aligned on a marker-by-marker basis using MAFFT local pair alignment (settings: max iterations = 1000; ep = 0.123; op = 3; --adjust-direction). We screened each alignment for samples ≥40% divergent from consensus sequences, which were almost always incorrectly assigned contigs. Alignments were retained if they included four or more taxa, had ≥100 bp length, and mean sample specificities (i.e., the “breadth of coverage” of the sample; see below) ≥50% across the alignment (to prevent non-overlapping segments of the alignment). We then separated alignments into two initial datasets: (1) “Exons-Only,” which included only exon contigs with intronic region trimmed from each alignment using the *Nanorana* genome sequence reference exon as a guide; and (2) “All-Markers,” which included the entire matching contig to the reference marker, including UCE markers. These two sets of alignments were only externally trimmed, using a custom R script, resulting in alignments in which at least 50% of the samples have sequence data at the both alignment ends. These alignments were not internally trimmed and should not be used until further processing (see below).

#### Final Sequence Alignments

The Exons-Only and All-Markers alignment sets were further trimmed into usable phylogenetic analyses datasets, and data type comparisons, resulting in: (1) “Exons,” each exons-only alignment was adjusted to be in an open-reading frame in multiples of three bases and trimmed to the largest reading frame that accommodated >90% of the sequences; (2) “Introns,” a consensus sequence of the exon previously delimited was aligned to the All-Markers dataset and the aligning region was removed leaving only the two intron ends, which were concatenated; (3) “UCEs,” were separately saved and not modified; (4) “Legacy,” markers from prior studies (excluding UCEs) were saved separately for ease of access and comparison; (5) “Gene,” exons from the dataset above were concatenated and grouped together if they were found from the same predicted gene from both the *Nanorana* and *Xenopus* genomes because these exons are likely to be strongly linked genetically (the exons could also be unlinked because of long introns so all datasets should be considered). Finally, the Introns and UCE datasets were internally trimmed using trimAl (automatic1 function; Capella-Gutiérrez et al. 2009). Our five trimmed final datasets were formatted as Phylip files and are suitable for phylogenetic analyses.

### Sequence capture evaluation

#### Sequence capture sensitivity

We evaluated the “sensitivity” of the sequence capture results, where sensitivity (also termed “breadth of coverage”) is defined as the percent of bases of a target marker covered by a post-assembly contig. To calculate sensitivity, we used the collection of target markers from probe design within the Hyloidea (n=24 samples), Ranoidea (24), and Reduced-Ranoidea (30) probe set samples and compared them to the matching contig for each of the sequenced samples using BLAST (*dc-megablast*). We did not evaluate introns, because they were not specifically targeted, but assessed their length separately. After matching with BLAST, we calculated the percent sensitivity per target marker and each sample by dividing the total base-pair length of the target marker sequence by the length of the matching portions of the sample contig.

#### Sequence capture specificity

“Specificity” refers to the percentage of cleaned reads (from the “cleaned-reads” set) that can be mapped back to the target markers. We assessed specificity within the Hyloidea (n=24 samples), Ranoidea (24), and Reduced-Ranoidea (30) probe set samples. First, we created a reference from the markers targeted with the probe set and mapped cleaned reads from each sample to the target markers using the program BWA v0.712 (Li and Durbin 2009). Each reference was indexed (function: *bwa index*) and reads from each sample were mapped to a reference (function: *bwa mem*), using SAMTOOLS (Li et al. 2009) to convert between file-types (functions: *view* and *fastq*). Finally, we counted the number of cleaned reads mapping back to the reference markers, to calculate the specificity (number mapped reads / total cleaned reads).

#### Sequence capture missing data

To assess levels of missing data from the different probe sets, we characterized variation in two ways: (1) missing data at the base pair level refers to targeted base pairs that are missing from a sample in an alignment (called “missing base pair data” throughout; also the inverse of sensitivity from above); and (2) missing data at the marker level (called “missing marker data” throughout) refers to targeted markers that were not adequately captured for a sample after applying the standard alignment filtering from above. To calculate missing base pair data, we counted the number of base pairs for each sample for each alignment and divided by the total length of the alignment. To calculate missing marker data, we used the final All-Markers alignments after the post-processing filtration and trimming steps described above and calculated missing marker data as the percent of markers not included in the final set of alignments for each sample.

#### Effect of genetic distance

To predict whether phylogenetic distance impacts sensitivity, specificity, and missing marker data, rather than stochastic factors or sequencing effort, we test for a linear relationship between these calculated variables and pairwise genetic distance from the design markers. We compared the genome-designed Ranoidea probe set and consensus-based sequence design from the Hyloidea probe set. Genetic distance was calculated using an uncorrected pairwise distance and the mean was computed across markers for each sample; for Ranoidea, this distance was calculated from the *Nanorana* genome sequence used in creating probe sequences, whereas in Hyloidea, distance was calculated from the consensus sequences created for the target markers using available sequence from the Hyloidea transcriptomes. For these analyses, we only included markers shared between by both probe sets (totaling 6,130 markers). A significant negative relationship would suggest that lower sensitivity and specificity is driven by higher sample genetic distance from the design markers. A significant positive relationship between genetic distance and missing marker data would suggest that sample dissimilarity from the design markers leads to missing marker data.

#### Marker depth of coverage

The “depth of coverage” or “depth” for targeted sequence capture data was calculated, where depth refers to the number of bases from the cleaned reads that overlap a given assembled base or bin of bases from each of the target markers (often denoted as “X”). We first created a reference for each samples’ set of post-assembly contigs that were targeted with the probe set and next mapped cleaned reads to these contigs using BWA (“bwa-mem” function). Next we removed exact duplicate reads using PICARD TOOLS (http://broadinstitute.github.io/picard/). To calculate a per-base overlap of cleaned reads to contig base pairs, the ‘depth’ function was used from SAMTOOLS. Depth was calculated across all targeted markers and samples for every base pair and was binned into 1% sized bins.

To describe the variation in depth among samples and markers separately, we calculated two metrics: (1) sample depth, median depth of markers calculated for each sample; and (2) marker depth, for each marker, median depth of samples calculated for that marker. We used median values, because individual depth measurements are not centered on zero and have a positive skew, such that few samples / markers have extremely high depth values, biasing towards much higher mean values. We calculated the marker depth separately for exons and introns, across markers. We used a Student’s two-sample *t*-test to test for a significant difference between depth of coverage from our Reduced-Ranoidea 20K set and Ranoidea 40K sets. We transformed values to fit a normal distribution and, if that assumption could not be met, we used a Mann-Whitney U test, which does not assume a normal distribution. Statistical tests were performed in R.

#### Phylogenetic scale

We evaluated each of our All-Marker, Exon, Intron, UCE and Gene datasets for criteria typically considered informative in modern, model-based phylogenetic inference (e.g. Townsend 2007). We calculated statistics for each marker, including: number of taxa, alignment length (total bp), percentage of missing base pair data (percent of missing bases across alignments), percentage of missing marker data (number of missing taxa from alignments), number of informative sites, and percent of informative sites. Additionally, we addressed whether Exon alignment length is related to the number of informative sites, by testing for a positive relationship between these two variables using trimmed datasets for each phylogenetic scale. These metrics were calculated using the Alignment Assessment tool from Portik et al. (2016).

#### Population genetics

To test whether these data could be used for population genetics, we aimed to locate variants and SNPs that are high quality and have sufficient depth of coverage. We note that these statistics depend entirely on the output from the sequencing platform being used, number of samples multiplexed per lane, and number of markers from the probe set included during hybridization enrichment; therefore it would be difficult to compare results across different study designs and we demonstrate possible results if researchers follow our design. We used GATK v4.1 (McKenna et al. 2010), following developer best practices recommendations for discovering and calling variants (Van der Auwera et al. 2011).

To discover potential variant data (e.g. SNPs, indels), we used a consensus sequence from each alignment from the target group as a reference and mapped cleaned reads back to reference markers from each sample. We used BWA (“bwa mem” function) to map cleaned reads (cleaned-reads dataset explained above) to our reference markers, adding the read group information (e.g. Flowcell, Lane, Library) obtained from the fastq header files. Next we used SAMTOOLS to convert the mapped reads SAM file to a cleaned BAM file, and merged BAM files with our unmapped reads, as required for downstream analyses. We used the program PICARD to mark exact duplicate reads that may have resulted from optical and PCR artifacts, and reformatted each dataset for variant calling. To locate variant and invariant sites, we used GATK4 to generate a preliminary variant dataset using the GATK program *HaplotypeCaller,* to call haplotypes, in GVCF format, for each sample individually.

After processing each sample, we used the GATK *GenomicsDBImport* program to aggregate samples from separate datasets into their own combined database. Using these databases, we used the *GenotypeGVCF* function to genotype the combined sample datasets and output separate “.vcf” files for each marker, containing variant data, from all samples, for final filtration. The preliminary variant set was filtered into four datasets: (1) All variants, variants kept after moderate filtering to remove probable errors filtered at a quality score > 5; (2) High quality variants, variants included SNPs, MNPs, and indels filtered at a quality > 20; (3) SNPs, the number of SNPs after high-quality filtering (quality > 20); and (4) unique markers, the number of unique markers from the probe sets that contain at least one high quality SNP.

## Results

### Sequence capture evaluation

We sequenced 105 samples, resulting in a mean base pair yield of 1,234 ± 577 mega base pairs (Mbp; range: 466–4,321 Mb) and a mean 8,172,461 ± 3,827,207 (range: 3,090,636–28,621,450) paired reads for each sample (Figure 2; for raw results per sample see Supplementary Table S4 and Supplementary Figure S1). Filtering raw reads to remove exact duplicates, low complexity and poor-quality bases, adapter and contamination from other non-target organisms resulted in a mean 84.5% ± 11% of reads (range: 27–96%) passing the quality filtration steps (mean: 1,041 ± 544 Mbp; range: 233–3,900 Mbp). After merging paired-end reads and reducing redundant reads, we recovered a mean of 443,032 ± 275,084 (range: 159,958–2,123,190), merged paired-end reads and singletons were used as input for assembly. After assembly, our samples yielded a mean of 15,832 ± 5,575.9 (range: 6,968–43,113) contigs, with a mean length of 860 ± 92 (range: 128–24,355) base pairs (Figure 2).

**Figure 2.**
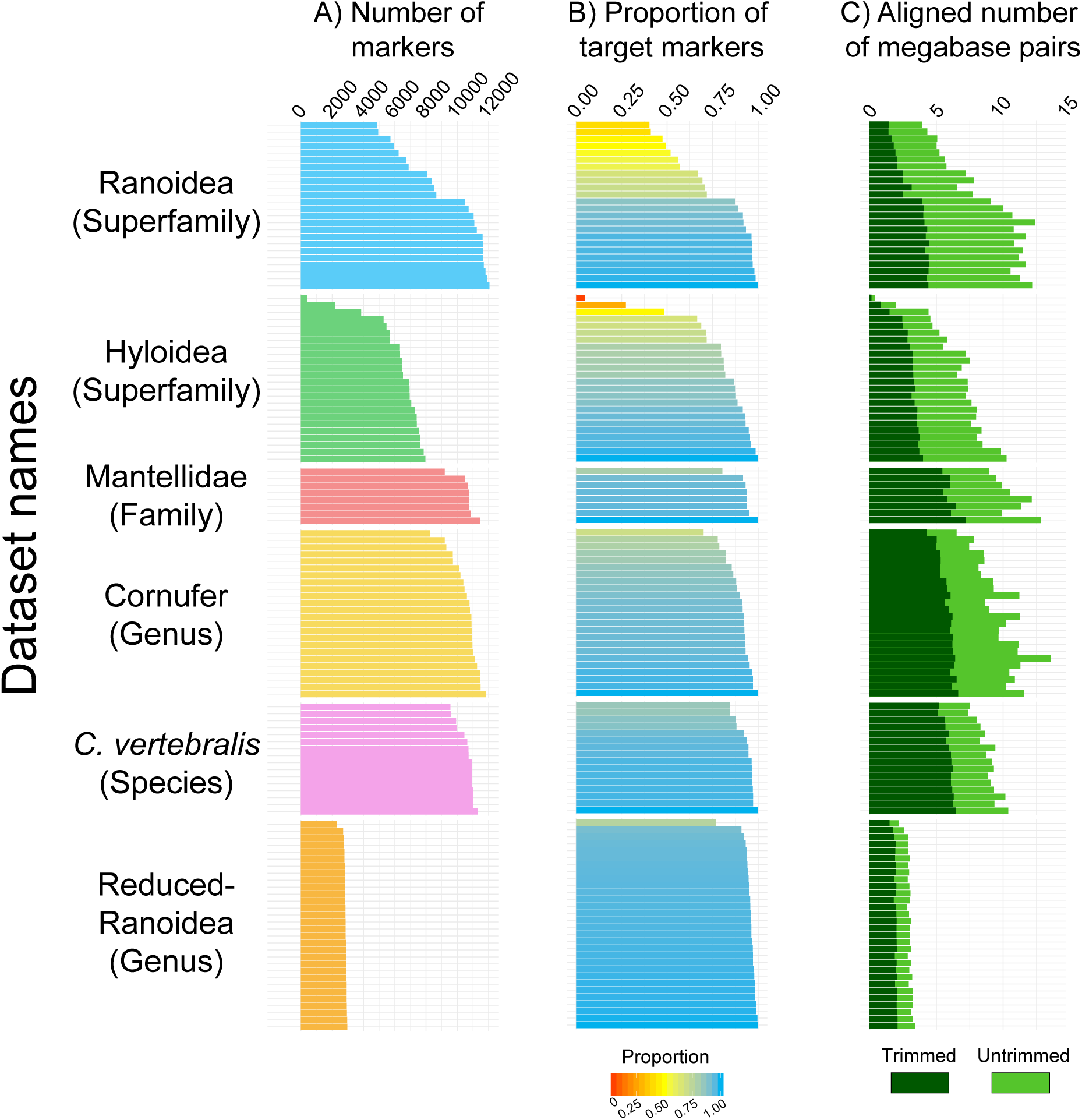
A summary of the data obtained for each sample, shown at each phylogenetic scale. The total number of markers in (A) are those obtained after matching to the target markers and removal of paralogs. In (B), the proportion is the sample total number of markers divided by the number of target markers. Finally, in (C) the total number of megabase pairs are the number of aligned bases before (light green) and after trimming (dark green). Note that the Hyloidea probe set sampling included one Salamander and two Archaeobatrachian frogs (the first 3 samples), which were tested and performed poorly compared to the Hyloidea clade samples. More detailed per sample information from this figure can be found in Supplementary Figure S1.

**Table 4.** Summary of the All-Markers dataset for each phylogenetic scale. The abbreviation PIS refers to parsimony informative sites.

| Clade<br>(Scale) | Dataset | N samples<br>(mean ± sd) | N<br>markers | Total bp | Length | Total PIS | PIS<br>(mean ± sd) |
| --- | --- | --- | --- | --- | --- | --- | --- |
| Anura<br>(Order) | exon | 34.3 ± 7.9 | 4566 | 961,305 | 210.5 ± 145.4 | 409,774 | 40.4 ± 8.9 |
| | intron | 31.6 ± 7.9 | 4185 | 1,317,305 | 314.8 ± 171.4 | 1,041,290 | 80.5 ± 14.2 |
| | legacy | 42.8 ± 3.5 | 15 | 15,258 | 1017.2 ± 412.2 | 5,523 | 35.6 ± 6.7 |
| | gene | 41.6 ± 4.7 | 857 | 457,743 | 534.1 ± 323.7 | 190,521 | 40.6 ± 6.4 |
| | UCE | 42.3 ± 5.7 | 567 | 319,259 | 563.1 ± 178.8 | 128,088 | 39.3 ± 12.3 |
| Hyloidea<br>(Superfamily) | exon | 16.2 ± 5.2 | 5696 | 1,332,048 | 233.9 ± 241.3 | 307,447 | 20.9 ± 8.6 |
| | intron | 15.6 ± 4.9 | 5636 | 2,310,925 | 410 ± 199.1 | 1,174,156 | 52.5 ± 20.3 |
| | legacy | 21.1 ± 4.1 | 84 | 79,113 | 941.8 ± 321.6 | 19,900 | 25.2 ± 6.7 |
| | gene | 19.9 ± 3.1 | 985 | 546,951 | 555.3 ± 376.6 | 121,821 | 21.1 ± 6.1 |
| | UCE | 20.9 ± 3.2 | 2359 | 1,941,867 | 823.2 ± 232.2 | 519,351 | 26.5 ± 10.3 |
| Ranoidea<br>(Superfamily) | exon | 17 ± 5 | 12149 | 2,453,364 | 201.9 ± 149.7 | 718,950 | 27 ± 9.2 |
| | intron | 16.1 ± 4.4 | 2982 | 1,104,972 | 370.5 ± 134.7 | 673,284 | 62.4 ± 16.6 |
| | legacy | 20.6 ± 4.4 | 24 | 21,186 | 882.8 ± 415.1 | 5,162 | 23.2 ± 7.5 |
| | gene | 21.3 ± 3 | 2235 | 1,347,147 | 602.8 ± 417 | 388,411 | 27.6 ± 6.5 |
| | UCE | 20.6 ± 4.7 | 650 | 442,500 | 680.8 ± 173.8 | 138,803 | 31 ± 10 |
| Mantellidae<br>(Family) | exon | 7 ± 1.1 | 11339 | 2,413,164 | 212.8 ± 190 | 135,430 | 5.2 ± 2.8 |
| | intron | 7 ± 1.1 | 11314 | 6,700,598 | 592.2 ± 206.8 | 1,050,533 | 16 ± 7.7 |
| | legacy | 7.5 ± 1.1 | 29 | 35,352 | 1219 ± 787 | 1,621 | 4.4 ± 2.2 |
| | gene | 7.8 ± 0.5 | 2077 | 1,275,423 | 614.1 ± 445.5 | 68,487 | 5.2 ± 2 |
| | UCE | 7.4 ± 1 | 604 | 545,819 | 903.7 ± 215.8 | 32,938 | 5.9 ± 3.1 |
| *Cornufer*<br>(Genus) | exon | 20 ± 4.8 | 11990 | 2,609,055 | 217.6 ± 231.3 | 177,481 | 6.2 ± 3.4 |
| | intron | 19.9 ± 4.7 | 11967 | 6,046,801 | 505.3 ± 159.8 | 1,108,147 | 18.4 ± 7.8 |
| | legacy | 22.1 ± 4 | 29 | 36,594 | 1261.9 ± 828.8 | 2,198 | 5.8 ± 3.7 |
| | gene | 23.2 ± 1.8 | 2203 | 1,368,702 | 621.3 ± 459.2 | 85,886 | 6.1 ± 2.3 |
| | UCE | 20.6 ± 5 | 628 | 473,984 | 754.8 ± 167 | 31,250 | 6.4 ± 4.1 |
| *Cornufer*<br>*vertebralis*<br>(Species) | exon | 14.1 ± 2.9 | 11393 | 2,507,571 | 220.1 ± 207 | 18,347 | 0.7 ± 1.2 |
| | intron | 14.1 ± 2.9 | 11375 | 5,790,769 | 509.1 ± 134.8 | 189,974 | 3.3 ± 4.1 |
| | legacy | 14.6 ± 2.7 | 29 | 36,036 | 1242.6 ± 826.9 | 201 | 0.5 ± 0.4 |
| | gene | 15.7 ± 1 | 2100 | 1,314,084 | 625.8 ± 460.3 | 8,657 | 0.7 ± 0.7 |
| | UCE | 14.5 ± 2.7 | 602 | 436,702 | 725.4 ± 127.5 | 4,564 | 1 ± 1.8 |
| *Occidozyga*<br>(Reduced) | exon | 27 ± 5.3 | 3110 | 1292427 | 415.6 ± 678.3 | 75,773 | 5.3 ± 2.8 |
| | intron | 26.8 ± 5.3 | 3100 | 1472238 | 474.9 ± 138.5 | 235,791 | 16 ± 7.1 |
| | legacy | 28 ± 5.9 | 85 | 88188 | 1037.5 ± 551.9 | 3,842 | 4.2 ± 2.3 |
| | gene | 29.6 ± 1.9 | 213 | 156186 | 733.3 ± 718.1 | 8,216 | 5.1 ± 2.3 |
| | UCE | 24.8 ± 10.5 | 4 | 3662 | 915.5 ± 311.1 | 260 | 6.6 ± 6 |

The assembled contigs from each sample were next matched using BLAST to the target markers from each of the three probe sets (Hyloidea, Ranoidea, Reduced-Ranoidea). The Hyloidea set produced a mean of 7,443 ± 2,022 (range: 616–9,570) contigs that were matched uniquely to target markers (mean marker proportion: 0.701 ± 0.199; range: 0.058–0.901). The Ranoidea set produced a mean of 10,304 ± 1,645 (range: 5,050–12,235) contigs that matched uniquely to target marker set (mean marker proportion: 0.757 ± 0.121; range: 0.371–0.898). Finally, The Reduced-Ranoidea set produced a mean of 2,847 ± 123 (range: 2,299–2,983) unique contig matches (mean marker proportion: 0.877 ± 0.038; range: 0.708–0.919).

### Sequence capture sensitivity

Sequence capture “sensitivity” was measured across our three probe sets (Hyloidea, Ranoidea, Reduced-Ranoidea) by assessing the percent of bases of target markers (exons and UCEs) covered by post-assembly contigs. The mean sensitivity across exons from samples in the Hyloidea set was 70.3 ± 20% (*n* = 24; range: 5.7–89.9%); the Ranoidea set mean sensitivity was 70.6 ± 19% (*n* = 24; range: 36.9–93.1%); and our Reduced-Ranoidea set mean sensitivity was 89.1 ± 4 % (*n* = 30; range: 71.6–93.5%; Table 2; Figure 3A). For UCEs, the mean sensitivity from samples in the Hyloidea set was 87.1% ± 16.9% (*n* = 24; range: 15.5– 96.6%), the Ranoidea set was 85.7 ± 8% (*n* = 24; range: 62.2–94.6%), and our Reduced-Ranoidea set did not include UCEs.

**Figure 3.**
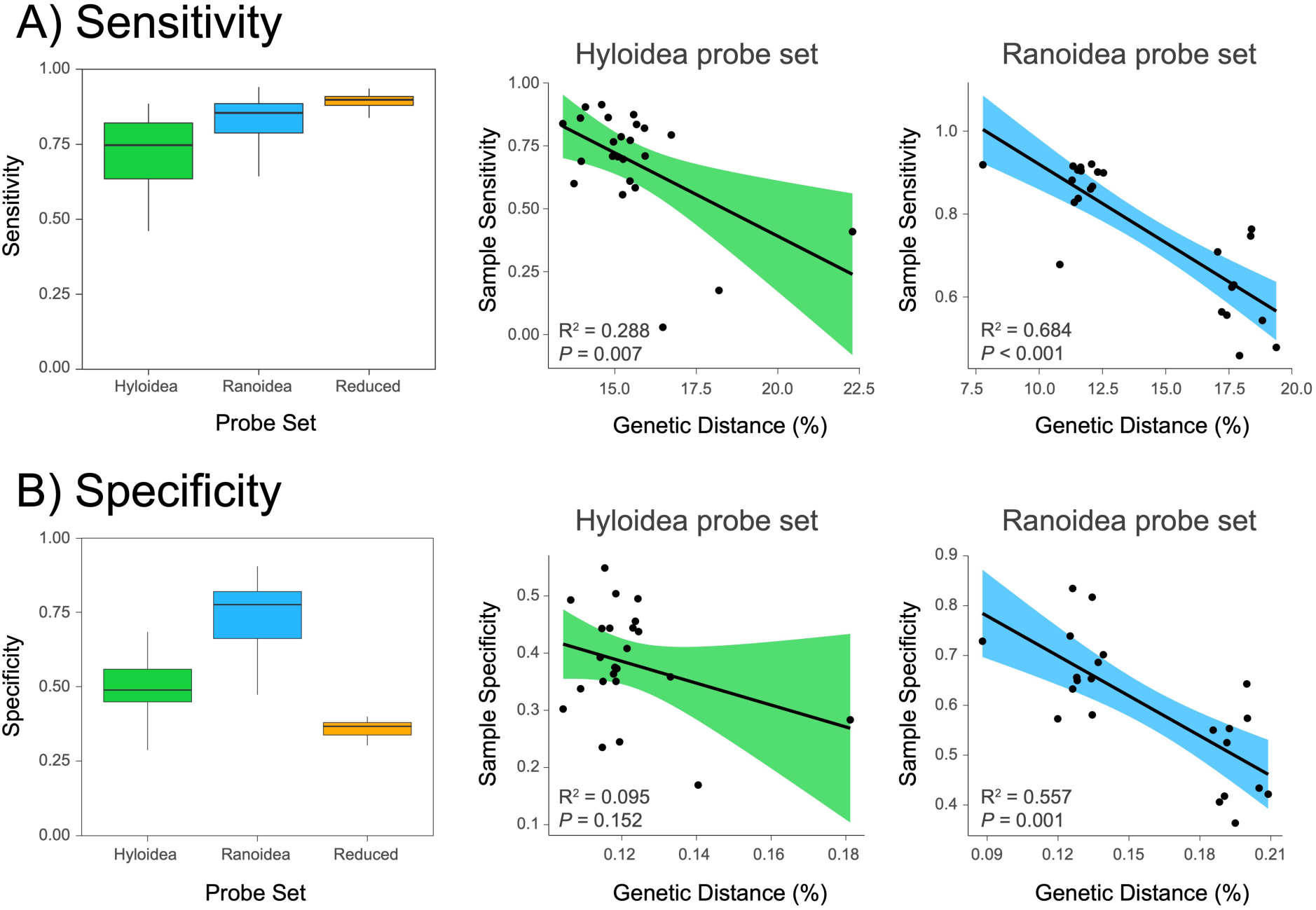
Sample sensitivity and specificity compared among three different probe sets (Hyloidea, Ranoidea, and Reduced-Ranoidea). In addition, relationships between genetic distance and sensitivity / specificity, evaluated for Hyloidea and Ranoidea probe sets (Reduced-Ranoidea not included because samples are all from a single genus and have the same genetic distances). The box plots show the distribution of sensitivity (A) and specificity (B) values across the probe sets. Sensitivity (A) has a significant negative relationship with genetic distance in Ranoidea and a weaker but still significant relationship in Hyloidea. Specificity (B) has a significant negative relationship in Ranoidea and a weak and non-significant relationship in Hyloidea. The 95 percent confidence intervals of the estimated regression line are indicated by green and blue shading.

### Sequence capture specificity

Specificity (the percentage of cleaned reads that can be mapped back to the target markers) was assessed within our Hyloidea, Ranoidea, and Reduced-Ranoidea probe sets. The Hyloidea set had a mean specificity of 47.9 ± 13.8% (*n* = 24; range: 1.7–68.3%); our Ranoidea set, had a mean specificity was substantially higher at 71.1 ± 12.8% (*n* = 24; range: 47.3–90.5%); and surprisingly, specificity was lowest in our Reduce-Ranoidea set, with a mean of 35.7 ± 3.3% (*n* = 30; range: 24.5–40.0%; Table 2; Figure 3B).

### Sequence capture missing data

We assessed the amount of missing data for the three probe sets (Hyloidea, Ranoidea, Reduced-Ranoidea) by calculating the percent of missing base pairs (mbp) and missing markers (mm) out of the alignment lengths and number of target markers for each sample, respectively (Figure 4A–B). The mean mbp from samples in the Hyloidea set was 29.7 ± 20 % (*n* = 24; range: 10.1–94.3%), with a much larger range and variation because of the inclusion of divergent clades (Archaeobatrachian frogs and Salamanders). The Ranoidea set mean mbp was 29.4 ± 19 % (*n* = 24; range: 6.9–63.1%). Last, our Reduced-Ranoidea set mean mbp was 10.9 ± 4 % (*n* = 30; range: 6.5–28.4%; Table 2; Figure 4A). The Hyloidea set had a mean of 21.9 ± 22.7 % mm (*n* = 24; range: 0–95.0%), with a much larger range and variation from the divergent clades. In our Ranoidea set, mean mm was slightly higher at 22.7 ± 21.2 % (*n* = 24; range: 0–59.7%). Finally, mm in our Reduce-Ranoidea set was substantially lower, with a mean of 4.5 ± 4.2 % (*n* = 30; range: 0–23.2%; Table 2; Figure 4B).

**Figure 4.**
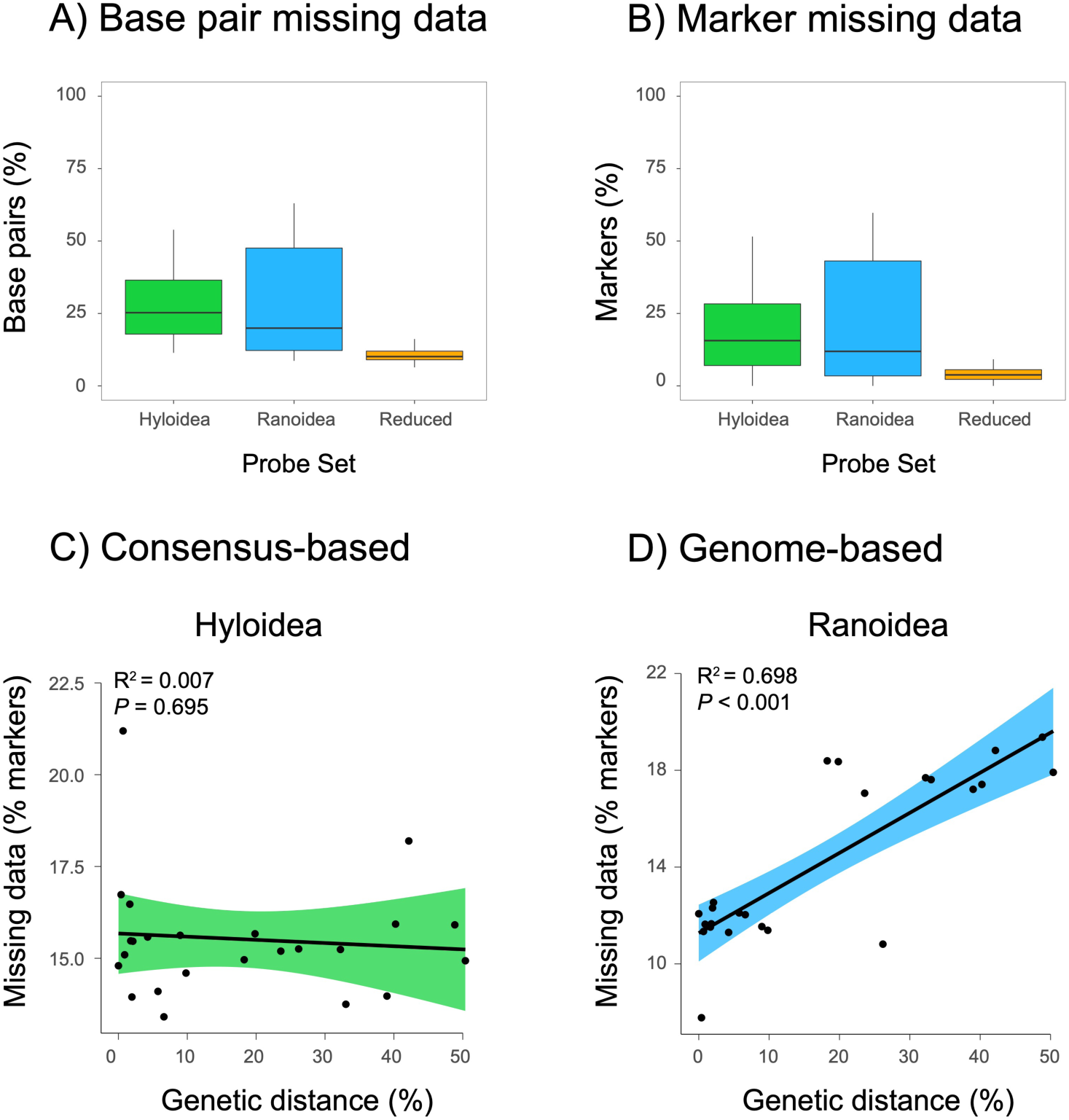
Missing data are compared for three different probe sets (Hyloidea, Ranoidea, Reduced-Ranoidea). Base pair missing data (also the inverse of sensitivity) in (A) is the percent of missing data calculated from the number of missing bases pairs across alignments for each sample. Marker missing data in (B) is the percent of missing markers across alignments for each sample. The consensus-based marker design missing data from Hyloidea in (C) has a weak and non-significant relationship with genetic distance. Conversely, the genome-based missing data from Ranoidea in (D) has a strong and significant positive relationship with genetic distance. Reduced-Ranoidea not included because samples are all from a single genus and have the same genetic distances). The 95 percent confidence intervals of the estimated regression line indicated with green and blue shading.

### Effect of genetic distance

To explore the impact of the genome-based Ranoidea and consensus sequences from transcriptomes in the Hyloidea probe sequences might lead to different sensitivity, specificity and missing data in sequence capture, we used linear regression to test for a relationship between these variables and genetic distance in our Ranoidea and Hyloidea datasets. For these tests, we calculated these measures from markers shared between both probe set datasets. The Reduced-Ranoidea samples were not tested because all of the samples are from the same genus with very similar genetic distances; therefore, phylogenetic distance from the design sequences would not be relevant.

When testing for sensitivity in the genome-designed Ranoidea samples, we found a strong negative relationship (*R*^2^ = 0.684; *P* < 0.001; Figure 3A), as described by the equation [sensitivity = 129.8 + −3.78 * average pairwise divergence], with pairwise divergence and sensitivity measured as percentages. Sensitivity decreased 3.78 percent for each percent increase of pairwise divergence. In addition, we found a weak but significant relationship among Hyloidea consensus sequence samples (*R*^2^=0.288; *P* = 0.007; Figure 3A), which was likely driven by three outlier data points (Figure 3A). In Hyloidea, sensitivity is equal to the equation [percent sensitivity = 171.5 + −6.62 * average percent pairwise divergence] where sensitivity decreased 6.62 percent for each percent increase of pairwise divergence.

To explore whether specificity is related to phylogenetic distance, we found a strong negative relationship (*R*^2^ = 0.632; *P* < 0.001) among our genome-designed Ranoidea samples (Figure 3B), equal to the equation [specificity = 102.0 + −2.67 * average pairwise divergence], and again with pairwise divergence and specificity measured as percentages. Specificity decreased 2.67 percent for each percent increase of pairwise divergence. Conversely, we found a weak and non-significant relationship within the Hyloidea consensus sequence samples (*R*^2^=0.121; *P* = 0.095; Figure 3B).

Finally, to ascertain whether increased divergence from the target marker sequence leads to more missing marker data, we tested for a relationship between percent of missing markers and genetic distance and found a strong positive relationship (*R*^2^ = 0.698; *P* < 0.001) in our genome-designed Ranoidea samples (Figure 4C). Missing marker data in Ranoidea is equal to the equation [missing data = 11.27 + 1.66 * average pairwise distance], when pairwise distance and missing data are measured as percentages. Missing marker data increased 1.66 percent for each percent increase of pairwise divergence. In the consensus-designed Hyloidea samples, we found a weak and non-significant relationship (*R*^2^=0.007; *P* = 0.695; Figure 4D).

### Marker depth of coverage

We assessed depth of coverage across markers and, also, separately for exons and introns (from the post-assembly contigs not alignments), for exons targeted with probes, and introns that were incidentally sequenced. Depth was calculated from the median number of cleaned-read base pairs that overlapped with the sample sequenced region per 1 percent bin. We compared depth using three different metrics: (1) we calculated sample depth (median depth of all markers for each sample; Figure 5A); (2) marker depth (median depth of all samples for each marker; Figure 5B); and (3) marker depth for the exon and intron, separately, from all samples, for each marker (Figure 5C).

**Figure 5.**
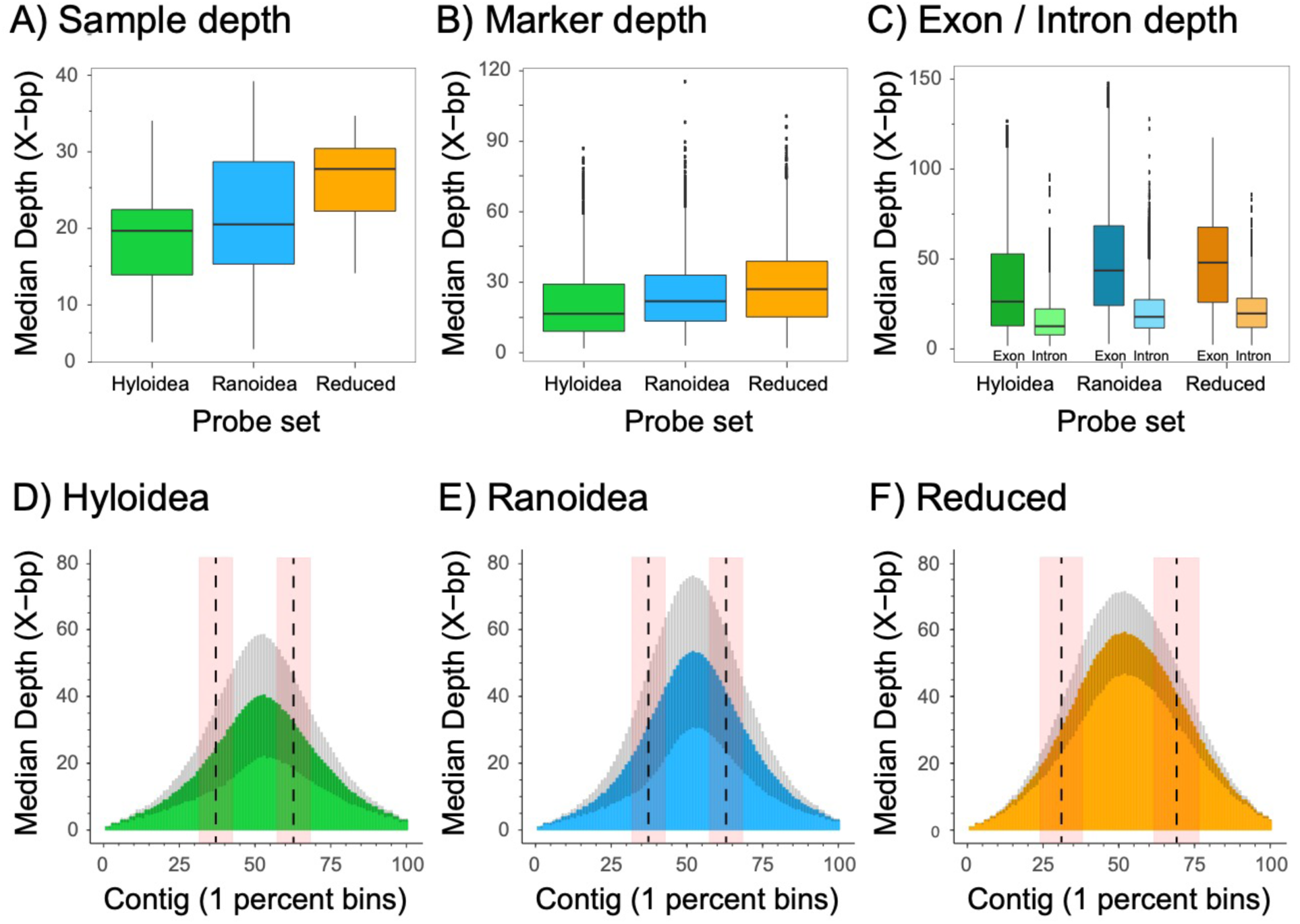
Depth of coverage statistics calculated for three different probe sets. Depth is summarized for A) median depth of markers within samples; B) median depth calculated between markers; C) median depth of markers was calculated for the exon and intron separately; and (D–F). Depth was calculated across each full-length marker contig by aligning the cleaned reads to the aligned sample contigs and counting the number of bases within each 1 percent bin (100 total bins per marker), and the median for each bin across markers was plotted. The gray coloration represents the standard deviation within each 1 percent bin across markers. The vertical dotted lines in each plot give the mean position where exon-intron boundaries occur, with the standard deviation shown in the red-colored shading.

In our Hyloidea probe set, we found a median sample depth of 19.7 ± 6.9 X (*n* = 24; range: 5.1–34.1 X; Figure 5A) and in Ranoidea, we found a median sample depth of 22.1 ± 8.5 X (*n* = 24; range: 4.2–39.2 X; Figure 5A). Finally, our Reduced-Ranoidea probe set had a median sample depth of 26.8 ± 5.5 X (*n* = 30; range: 14.2–34.7 X; Figure 5A). Because of small sample size and non-normal distributions, we used a Mann-Whitney U test and found a significant difference in the mean ranks of the median sample depth between the Reduced-Ranoidea 20K and Ranoidea 40K probe sets (*U* = 236; *P* = 0.031).

We assessed the median depth across markers, and found the Hyloidea probe had a median marker depth of 16.6 ± 13.5 X (*n* = 24; range: 1.8–87.0 X; Figure 5B, D) and the Ranoidea probe set had a median marker depth of 21.9 ± 14.0 X (*n* = 24; range: 3–115 X; Figure 5B, E). The Reduced-Ranoidea probe set had a median marker depth of 27.1 ± 16.0 X (*n* = 30; range: 2–100.7 X; Figure 5B, F), which was higher than the Ranoidea 40K set. Using a Student’s two-sample *t*-test, we found a significant difference in the distribution of median depth of markers between the Reduced-Ranoidea 20K and the Ranoidea 40K probe set (*ln* transformed*; T* = −9.346; *df* = 4464.6; *P* < 0.001). It is possible that the unshared markers between the 20K and 40K sets led to this difference, so we test only markers shared between the two sets (3,054 markers); we still found a significant difference in depth of coverage between the median depth of markers (*ln* transformed; *T* = 4.8; *df* = 6105.2; *P* < 0.001). Unexpectedly, the 40K had a higher median depth than the 20K across shared markers, but the overall median difference was very small (Ranoidea = 26.9 X; Reduced = 26.5 X).

To evaluate exons and introns separately, we measured the same parameters as above across all the samples. When measuring depth in the Hyloidea dataset, we found a median depth of 40.4 ± 29.3 X (*n* = 24; range: 2.3–129.8 X) for exons and a reduced median depth of 13.8 ± 10.7 X (*n* = 24; range: 1.3–72.9 X) for introns (Figure 5C). In our Ranoidea dataset, we found a median exon depth of 43.6 ± 29.6 X (*n* = 24; range: 2.7–147.7 X) and a lesser median intron depth of 13.2 ± 9.7 X (*n* = 24; range: 1.5–96.2 X; Figure 5C). Finally, for our 20K Reduced-Ranoidea data, we found a median depth of 48.1 ± 25.7 X (*n* = 30; range: 2.3–117.6 X) for exons and a lesser median depth of 15.8 ± 8.9 X (*n* = 30; range: 1.4–64.6 X) for introns (Figure 5C). Within the whole contig (Figure 5D–F), depth is generally highest near the center of the sequence and decreased towards its two ends. When binning contigs in 1 percent bins, the distance from the either edge to the start of the exon is ∼32–37 percent of the contig itself (Figure 5D–F). Interestingly, only ∼25 percent of any given contig is targeted exon sequence, whereas ∼75 percent is intron sequence, demonstrating that incidental capture is 3X higher than targeted areas. These results also show that when considering the exon separately from the intron, the median depth between the 20K and 40K probe sets are nearly the same.

### Phylogenetic scale

The resulting number of alignments before delimiting into data types (Exons, Introns, UCEs, Legacy, and Genes; the All-Markers dataset) for each of the phylogenetic scales was variable (Table 3; Figure 6A–B). Our Reduced-Ranoidea probe set (genus: *Occidozyga*) had 3,156 alignments (out of 3,247 targeted markers) created from the reduced probe set of 20,020 baits. Among the larger 40K bait sets, the scale with the fewest alignments was the Order-level scale with 5,500 alignments, which were shared after filtration and trimming (compared to the original 6,130 markers shared across the Ranoidea and Hyloidea probe sets). Hyloidea resulted in 8,712 alignments (because this probe set intentionally was designed with longer alignments) and the remainder of the scales had similar numbers of alignments at 12,009–13,099 because they resulted from the Ranoidea set (Table 3). Counts of alignments and other statistics across data types across phylogenetic scales are shown in Table 4 and Figure 6.

**Figure 6.**
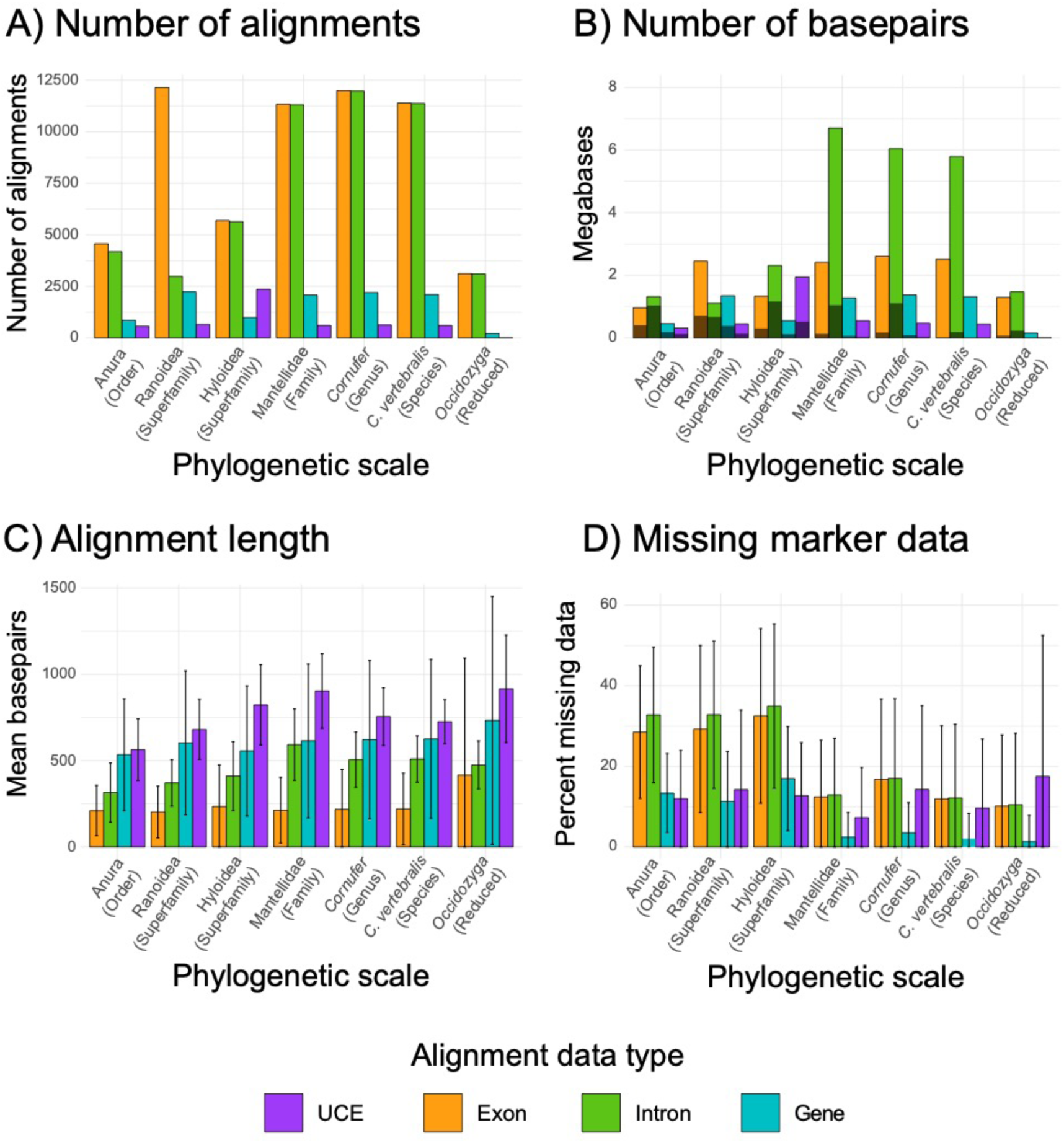
Summary statistics of alignments, comparing quantity of sequence data for different dataset types across different phylogenetic scales after filtration, alignment, trimming and dataset partitioning. Comparisons include: (A) number of alignments for each dataset at each phylogenetic scale; (B) number of base pairs and parsimony informative sites (shown as the darker shading) plotted for each dataset within each phylogenetic scale; (C) mean alignment length (with standard deviation shown as error bars) is compared for different data types and phylogenetic scales; and (D) percent of missing marker data (measured across markers with standard deviation shown as error bars) is compared for different datasets.

Mean alignment length was variable among the phylogenetic scales. Alignments at the Order-level scale were among the shortest, which can be largely attributed to trimming highly variable intron regions which were relatively more complex, resulting in more discarded alignments after filtration (Table 4; Figure 6C). Conversely, alignment lengths increased at more shallow phylogenetic scales, for which intron regions were less variable and easier to align due to less ambiguous inference of positional homology (Table 3). Missing marker data was also variable at various phylogenetic scales. In general, at the broadest phylogenetic scales (i.e. Order and Superfamily) there were higher levels of missing marker data (Figure 6D).

There was also a general pattern of all markers exhibiting increased variability (in terms of informative sites) at broader phylogenetic scales (Figure 5B; Figure 7). Not surprisingly, Introns were the most variable data type across all phylogenetic scales and even displayed at the species level (Table 4; Figure 7). Exons were moderately variable and similar to our Gene data type (which are composed of multiple exons). The UCEs were slightly less variable than the Exons and Genes, but still variation occurred in their characteristic flanking regions (admittedly, we selected the most variable UCEs for our probe sets). Finally, when testing whether alignment length is related to percentage of parsimony informative sites, we found a significant relationship across all phylogenetic scales. The strength of this relationship decreased at shallower phylogenetic scales (genus and species level), indicating marker variation generally decreased at shallower scales (Table 5). However, clear differences in variation exist between the Exons and Introns, with Introns providing a majority of the informative sites detected at shallower scales (Table 5; Figure 7).

**Figure 7.**
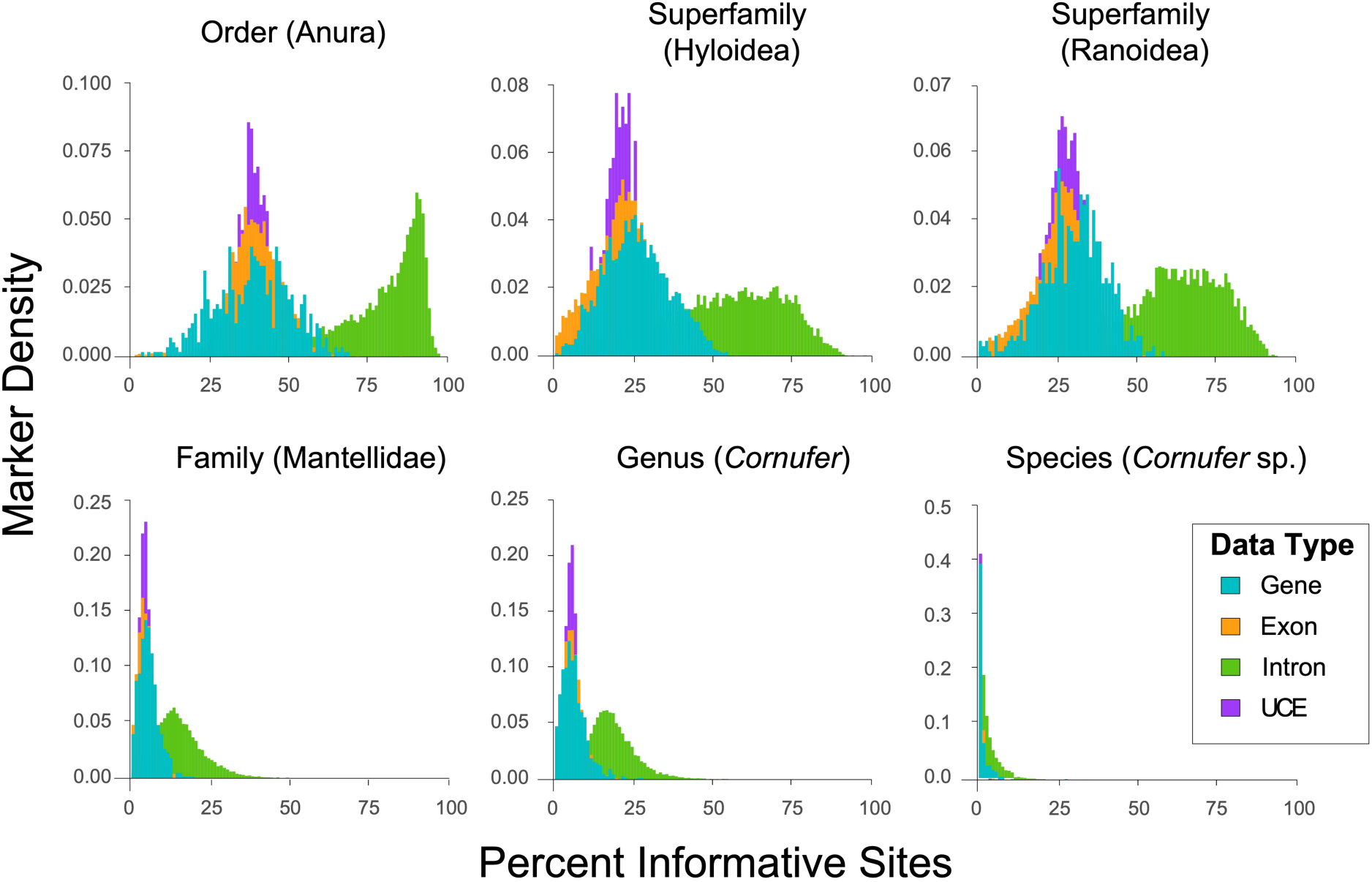
Comparisons of percent of parsimony informative sites, for each data type, at phylogenetic scales. Introns are the most variable data type across phylogenetic scales and display moderate variation even at the species level. Exons and UCEs had moderate amounts of variation, where both data types had little variation at the species level. The marker density has been normalized to best compare the different data types.

**Table 5.** Results from tests for a relationship between alignment length and parsimony informative sites across phylogenetic scales. The trimmed All-Markers dataset was used for this analysis.

| Scale | Clade | R<sup>2</sup> | P |
| --- | --- | --- | --- |
| Order | Anura | 0.799 | <0.001 |
| Superfamily | Ranoidea | 0.623 | <0.001 |
| Superfamily | Hyloidea | 0.523 | <0.001 |
| Family | Mantellidae | 0.286 | <0.001 |
| Genus | *Cornufer* | 0.289 | <0.001 |
| Species | *Cornufer vertebralis* | 0.030 | <0.001 |
| Reduced | *Occidozyga* | 0.445 | <0.001 |

### Population genetics

To assess whether FrogCap markers have utility in population genetic studies, we used GATK4 to discover genetic variants and SNPs across our sampling of taxonomic variation, within our seven datasets, addressing five phylogenetic scales. Generally, we found a pattern of decreasing variants (i.e. SNPs, indels) from higher to lower phylogenetic scales (Order to Species-levels), with greater than 20,000 variants at the species level after high quality filtering (Figure 8). This pattern remains for SNPs only, with greater than 10,000 SNPs found at the species level after aggressive filtering (Figure 8). Finally, in studies for which the independence of SNPs is required, we find at least one strongly supported SNP (and often numerous others) per each individual marker, thus permitting thousands of unlinked SNPs in FrogCap datasets, even at the species level (Figure 8).

**Figure 8.**
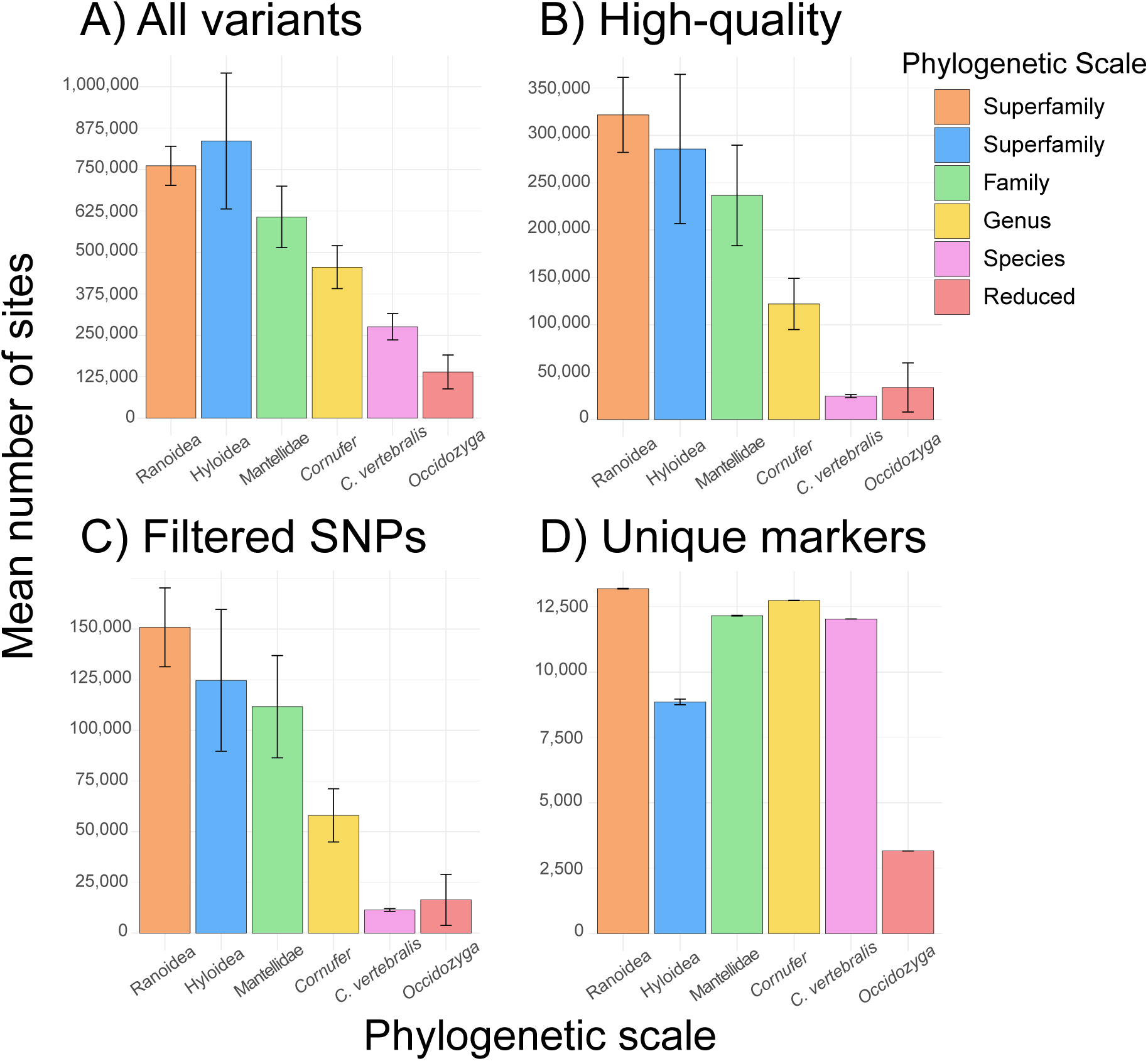
Mean number of variants across samples after different filtration schemes are shown at each phylogenetic scale (Anura not shown). The different variant filtering are: (A) all variants after moderate filtering (quality > 5); (B) High quality variants including SNPs, MNPs, and indels (quality > 20); (C) the number of SNPs after high-quality filtering (quality > 20); and (D) the number of unique markers from the probe sets that contain SNPs.

## Discussion

We introduce FrogCap, a publicly available collection of markers with associated sequence capture probe sets, representing a remarkably powerful suite of resources for the collection of genomic data for phylogenomics, systematics, and population genetics in any frog system. FrogCap is modular and topically-adaptable, providing a wealth of options for marker selection and analyses, which allows it to be tailored to specific study questions. FrogCap was created with a novel approach that identified orthologous markers from frog genomes and transcriptomes and includes two complementary probe sets (with 40,040 baits) designed to capture the same markers across major clades (Ranoidea and Hyloidea; >95% of frog diversity). We assessed three differently designed probe sets of FrogCap using 105 samples, selected strategically across five phylogenetic scales. Importantly, we demonstrate the success of FrogCap at each of these levels, suggesting our approach to sequence capture design can be applied to other groups of organisms. Based on the development of FrogCap and our sampling design, we are able to make several recommendations on key topics related to sequence capture. We discuss the effects of using different bait kit sizes, whether genome-based or consensus-based probe design are more successful, the impact of phylogenetic scale on sequence capture success, and the importance of incorporating different molecular marker types. Altogether these results suggest that the FrogCap marker set is adaptable and contains a variety of markers potentially useful for phylogenetic analyses and population genetics; this effort provide a much-needed novel mixture of molecular marker types for empirical comparison, side-by-side optimization, and exploration of data type combinations which have the potential to resolve difficult phylogenetic relationships.

### Bait kit size

One of the first decisions in sequence capture study design is selecting the size of the bait kit. Given the same library pooling design, sequencing effort, and probe tiling scheme, the number of probes available for capture design will directly affect the number of target markers and can indirectly affect the depth of coverage. Here we aimed for an optimal trade-off between depth of coverage and sequencing costs while maintaining the same conditions for multiplexing sample numbers and sequencing effort. Although larger sized bait kits can target more markers, they could be disadvantageous if the sequencing effort is not adequate, resulting in higher amounts of missing data. Additionally, a smaller bait kit that targets a subset of markers would be expected to have greater resulting depth per sample because fewer genomic areas are being targeted for sequencing. Given these expectations, we tested the effects of using a 20,020 and a 40,040 bait kit (20K and 40K respectively; MyBaits kits from Arbor Biosciences). To control for additional variables in this experiment, we used comparable library pooling designs, sequencing effort, and probe tiling schemes for each kit size. The Ranoidea and Reduced-Ranoidea sets were compared here because the Reduced-Ranoidea set uses a subset of the same bait sequences from the Ranoidea set.

Our results suggest that there might be an advantage to using the 20K bait set when compared to the 40K set, because using a smaller bait set resulted in a higher capture success. Specifically, we found the 20K bait set had a higher sensitivity (∼89% versus ∼70% in the 40K; Figure 3) and much fewer missing marker data overall (∼5% vs ∼22% in the 40K; Figure 4). When capture success is not related to sequencing effort, we would expect no relationship between genetic distance and sensitivity / missing data; here we found significant relationships between genetic distance of samples from design markers and missing marker data / sensitivity in the Ranoidea 40K bait set (Figure 3–4). These results suggest that the 40K set had adequate sequencing depth to sequence most of the markers, ruling out a potential advantage of the 20K set.

Despite the potential advantages regarding capture success and missing data, depth of coverage is another important factor to consider. Our expectation was for a 2X difference in depth between the 20K and 40K kits. Although depth of coverage was significantly different between the samples and markers in the kits, the difference was actually relatively small (medians: 40K = ∼24X; 20K = ∼26X; Figure 5). The same was true for a comparison of the shared markers only (medians: 20K = ∼26.5X; 40K = ∼26.9X). Although significant differences were found, both kits resulted in favorable depth of coverage values. Overall our results indicate that the 20K offers only a small advantage over the 40K, but we recommend using the 40K because it results in thousands more sequenced markers despite the increased capture success of the 20K. This ultimately results in more target markers recovered in the 40K despite the lowered capture success. Given the high depth of coverage found for the 20K kit in our experiment, it may be possible to multiplex more samples (possibly even the twice the number of samples), which could offer an advantage for projects focused on fewer markers and reduce costs.

### Probe design

There are numerous considerations for designing probes for sequence-capture experiments. Ultimately, the availability of genomic resources dictates how probes can be designed. The availability of a genome allows the detection of exon boundaries, inclusion of non-coding sequences (introns, UCEs), and enhanced paralog detection. Conversely, probe design based on transcriptomes is more challenging because paralog detection is difficult and the presence of intron-exon boundaries could interfere with capture success (but see Portik et al. 2016). Regarding both types of genomic resources, probe tiling density is an important factor. Higher densities result in many probes covering the target regions, which is expected to increase the likelihood of obtaining that region (i.e. AHE markers; Lemmon et al. 2012). In contrast, lower densities result in fewer probes per target, allowing more overall targets to be included but at the potential cost of lower capture success.

Our experiment included probes designed from a combination of transcriptomes and genomes. We found that the genome-based design (Ranoidea) had slightly more capture success than the transcriptome-based (Hyloidea) design. Many of the uncaptured target markers and paralogs detected post-processing in Hyloidea were transcriptome-based, whereas the shared markers between Ranoidea were largely successful. However, the transcriptome-based approach successfully recovered 70 percent of the target markers, which is consistent with other transcriptome-based studies (e.g. Bi et al. 2012; Portik et al. 2016; Reilly et al. 2019). Furthermore, the number of genomic resources available for designing the Ranoidea probe set was much higher (24 transcriptomes and 1 genome vs. 5 transcriptomes), and likely affected the sequence capture outcomes for Ranoidea and Hyloidea. For example, the Hyloidea transcriptome markers could not be filtered as rigorously as the Ranoidea markers, and the lack of phylogenetic diversity in sampling resulted in consensus probe sequences more similar to some groups in the clade.

Probe tiling density is another important variable that differentiates FrogCap from other frog probe sets. A higher tiling density limits the number of target markers but increases the likelihood of obtaining a target region; conversely, a lower tiling density allows for more target markers but may result in higher levels of missing data. In our study, we used a lower density 2X tiling scheme (at least 50 percent of each probe overlaps with adjacent probes) that has been recommended as a standard for sequence capture designs (Tewhey et al. 2009). We found a slightly lower capture success (i.e. specificity and missing data) than other studies with more dense tiling designs (specificity: this study = 70–75%; Portik et al. 2016 = 80%), which led to a higher amount of missing marker data (this study = 20–25%; Portik et al. 2016 = 8–10%). However, the Reduced-Ranoidea design had greater success than these densely tiled studies, indicating that careful selection of markers based on prior successful sequencing can actually circumvent limitations imposed by lower density tiling. Despite the increased capture success in the Reduced-Ranoidea and more densely tiled sequence capture designs, having more markers by using a 2X tiling scheme actually results in more markers being recovered (Figure 6). Therefore, we recommend less dense tiling schemes because the number of markers lost from the 2X tiling scheme is small compared to the number of markers gained from the higher number of markers being targeted.

### Genetic divergence from probes

Phylogenetic distance from the design markers is likely one of the most important and predictable factors affecting sequence capture performance (i.e. sensitivity, specificity, and missing marker data). Similar to other studies (Portik et al. 2016), we found higher genetic distances from a target lead to a decreased likelihood of capture for the Ranoidea probe set that used genome-based marker design. This probe set was designed using only the *Nanorana* genome sequences, rather than from multiple species or consensus sequences. Surprisingly, the consensus-sequence probe design for the Hyloidea set had poor or non-significant relationships between genetic distance and sequence capture performance metrics. This would suggest that other factors could explain missing markers, such as paralogs which are more prevalent in the transcriptome-based sequence capture samples and genetic distance for a subset of the samples not related to the design transcriptomes (discussed above). Overall, our results suggest that these markers designed using consensus sequences improved the sequence capture success when compared to genome-based (i.e. Ranoidea) or multispecies transcriptome-based (e.g., Portik et al. 2016) designs.

### Phylogenetic scale

Understanding how sequence capture experiments perform at different phylogenetic scales is important for decisions about marker selection and probe design. In our experiment, we assessed the impact of phylogenetic scale on the number of markers recovered, total base pairs recovered, the proportion of informative sites, and levels of missing data at the order, superfamily, family, genus and species level (Taylor & Piel 2004; Townsend 2007). Generally, at broader phylogenetic scales (e.g. order, superfamily) the number of alignments decreases because missing data were more prevalent (∼25%) as a result of marker drop-off in divergent samples (Figure 6D). At shallower phylogenetic scales (e.g. family, genus, species) missing data levels were lower (< 10%) and more uniform across samples (Figure 6D). At shallower scales, the number of markers recovered appears to be related to how divergent the target group is from the design probes (i.e. Microhylidae genus level comparison would differ from Ceratobatrachidae). To help remedy this issue, we designed the FrogCap probe sets are modular in design. For example, the Reduced-Ranoidea probe set was designed based on successful capture data from Ranoidea, where we selected markers that were successful and had longer alignment lengths. With this probe set, we find a higher capture success (sensitivity) and many fewer missing data across alignments (∼5%).

### Molecular marker type

We evaluated the properties of UCE, Exon, Intron and Gene (i.e. exons concatenated from the same gene; see **Materials and Methods**) sequence data within and across phylogenetic scales. UCEs and Exons were targeted in the marker design, and Gene alignments are exons combined from the same gene; however, intronic sequence was indirectly acquired as “by-catch” from by sequencing adjacent regions of a captured DNA fragment (e.g. Bi et al. 2012; Guo et al. 2012; Tewhey et al. 2015). Similar to Portik et al. (2016), the number of base pairs of data available from non-targeted intronic sequence was 2–3 times higher than explicitly targeted exon sequence (Figure 6), suggesting that intronic sequence is an abundant and potentially important resource in exon-capture. Additionally, intronic sequence could be valuable because of potential neutral evolution relative to exons which are typically functional and under selection, and UCEs which are likely under strong purifying selection to remain ultra-conserved (Halligan et al. 2004; Katzman et al. 2007; Stephen et al. 2008).

Perhaps the most important characteristic to evaluate across different data types is the informativeness or variability of the marker, which could potentially differ based on phylogenetic scale and data type. We show that parsimony informative sites vary markedly between both data type and phylogenetic scale (Figure 7). Among data types, the Intron is the most variable and is similar to UCEs in that the variability increases towards the flanks of the marker alignment; however, UCEs have moderate variability when including the central conserved region. Conversely, Exons have a small to moderate amount of variation. Introns show this pattern likely because of neutral evolution or weak selection (Halligan et al. 2008), whereas exons remain more conserved because they are under strong selection (Katzman et al. 2007). At shallow phylogenetic scales (species or genus) exons and UCEs have little variation, but Introns are quite variable (Figure 7). At broader phylogenetic scales (order, superfamily), Exons and UCEs are more variable. At these scales Introns become highly variable, and data could be lost due to the difficulty in aligning these regions.

The alignment length for exons in particular should be considered when including markers for analyses, and the FrogCap probe set includes a mixture of exons from different size classes (Figure 1B). We generally show that exon length is related to the number of informative sites, where longer exons provide more information for phylogenetic analyses (Table 5; Townsend 2007). The mean size of exons was ∼250 bp (Table 1) which is similar to the predicted mean exon length across the *Nanorana* and *Xenopus* genomes (∼200 bp). Short exons are advantageous because they would allow more SNPs to be incorporated into analyses that require genetically unlinked SNPs (Figure 8). The main disadvantage of short exons is that they may lack sufficient variability for strong support in phylogenetic analyses, which is important for summary species tree methods using individual gene trees. For these types of analyses, FrogCap also includes ∼1,000 long exons greater than 500 bp in length (and ∼500 exons greater than 1,000 bp; Table 1). Long exons would be ideal for analyses where the goal is to have fewer markers/partitions, but stronger statistical support among gene trees. The publicly available FrogCap pipeline and website (https://www.frogcap.com) separates alignments into these different data types so that researchers will have a great deal of flexibility in choosing their optimal marker attributes for individual strategies behind analyzing sequence data.

### Future directions

FrogCap is a new modular collection of markers and sequence capture probe sets for frogs, which forms an important foundation for future work. First, although not the focus of this work, a phylogenetic assessment of the different data types and how they perform at different phylogenetic scales across different types of analyses would be important for future FrogCap studies and other phylogenomic studies that include different data types (Hutter et al. *unpublished data*). Second, some smaller amphibian phylogenetic groups are underrepresented (Salamanders, Caecilians, and Archeaobatrachian frogs) in FrogCap; work is currently underway on a “universal” module of markers that could be captured reliably across amphibians (but at the cost of fewer informative sites). In addition, specialized subsets and additional probe sets including a greater number of markers have been designed for specific clades (the genus *Limnonectes*, and family Dendrobatidae), which emphasizes the modular, taxon-specific adaptability of FrogCap (*unpublished data*). Third, although sequence capture is not the best method for acquiring SNP data, we demonstrate that thousands of high-quality SNPs can be discovered; future studies will address the performance of SNP-based analyses resulting from FrogCap. Finally, we create a refined version of the Ranoidea probes set (Version 2.0) informed from the analyzed Ranoidea V1 in this study; the refined Ranoidea V2 excluded markers that did not capture well across our test samples; these were replaced with additional UCEs, Legacy markers, and longer exons. Both versions are available, along with Version 1 at https://github.com/chutter/FrogCap-Sequence-Capture.

## Data Availability

A major goal in disseminating these resources is to provide the probe set described and tested here as a freely available and open access resource. The probe set and other resources are licensed under a Creative Commons Attribution 3.0 United States (https://creativecommons.org/licenses/by/3.0/us/legalcode).

All raw sequencing reads are made available in the GenBank SRA (BioProject: available upon publication) upon acceptance of this manuscript in a peer-reviewed journal. All alignments analyzed and materials for replicating analyses are made available on the Open Science Framework (https://osf.io/gvbr5/) following manuscript publication. Finally, the data analysis pipeline and scripts for all analyses here, the probe set and marker files, and data for several outgroups for other researchers to use are now available on Carl R. Hutter’s GitHub (https://github.com/chutter/FrogCap-Sequence-Capture).

## Cost estimates

We provide a brief summary of cost estimates for this project, which are based off of current quotes from library preparation and sequencing providers we have worked with (as of October 2019). The cost of the probes amounts to ∼$25 per sample, library preparation (outsourced at Arbor Biosciences^TM^) is ∼$75 per sample, and sequencing is ∼$15 a sample ($1300 a lane at NovoGene^TM^), which totals $110 a sample. These costs were attained with a large sample volume (>192 samples). None of these costs are guaranteed and are likely to fluctuate, and we offer no guarantee of similar quotes from these providers. Additionally, we note that in-house library prep (rather than using a service) can substantially reduce the library prep costs, and purchasing more probes in bulk can reduce costs.

## Acknowledgements

For funding and financial support that made this research possible, we would like to thank KU Research Investment Council (RIC) Level II Award (No. 2300207) to RMB and R. G. Moyle, and KU Docking Fund support to RMB. We also acknowledge University of Kansas Graduate Studies support to CRH, University of Kansas Genome Sequencing Core to both CRH and RMB, USA National Science Foundation (Graduate Research Fellowship award numbers 1540502, 1451148, 0907996 to CRH), DEB systematic panel awards 1654388, 1557053, 0743491 to RMB; Doctoral Dissertation Improvement Grant 1701952 (to SLT), and 0804115 to C. D. Siler). Additional funding was provided by the American Philosophical Society, and the Society for the Study of Evolution. We thank the Malagasy authorities for approving research permits (N°298/13/MEF/SG/DGF/DCB.SAP/SCBSE, N°303/14/MEEMF/SG/DGF/DAPT/SCBT, N°329/15/MEEMF/SG/DGF/DAPT/SCBT); specimens were exported under permits: N°017N-EV01/MG14, N°055N-EA02/MG15, N°041N-EA01/MG16) and the organizations MICET and Centro ValBio for logistics assistance with permits and in the field. The Solomon Islands Forest Department approved field research protocols and export permits (to SLT, RMB, and R. G. Moyle), and the Philippine Department of Environment and Natural Resources (DENR) facilitated Memoranda of Agreement and Gratuitous Permits to Collect Biological Specimens, issued to RMB, 2005–2019). We also thank David Blackburn of the California Academy of Sciences, Chan Kin Onn of the National University of Singapore, and the Museum of Vertebrate Zoology at Berkeley for tissue samples. This paper constitutes contribution number XXX of the Auburn University Museum of Natural History for PLW. Finally, we thank Arbor BioSciences and their staff scientists for assistance in designing the probes and library preparation, especially Jake Enk and Alison Devault.

## Supplementary Material

Supplementary Table S1. GenBank accession numbers for transcriptomes and genomes for probe design.

Supplementary Table S2. Meta-data and sample information for each sample.

Supplementary Table S3. GenBank accessions numbers for the decontamination genome filtration step.

Supplementary Table S4. Per sample sequencing and processing statistics. Supplementary Figure S1. Detailed per sample summary statistics.

## Literature Cited

Alexander AM, Su Y-C, Oliveros CH, Olson KV, Travers SL, Brown RM. 2016. Genomic data reveals potential for hybridization, introgression, and incomplete lineage sorting to confound phylogenetic relationships in an adaptive radiation of narrow-mouth frogs. Evolution 71:475–488.

Alexander RP, Fang G, Rozowsky J, Snyder M, Gerstein MB. 2010. Annotating non-coding regions of the genome. Nat. Rev. Genet. 11:559–571.

AmphibiaWeb. 2019. Information on amphibian biology and conservation. Retrieved October 8, 2019, from http://www.amphibiaweb.org

Andermann T, Fernandes AM, Olsson U, Töpel M, Pfeil B, Oxelman B, Aleixo A, Faircloth BC, Antonelli A. 2019. Allele phasing greatly improves the phylogenetic utility of ultraconserved elements. Syst. Biol. 68:32–46.

Bankevich A, Nurk S, Antipov D, Gurevich AA, Dvorkin M, Kulikov AS, Lesin VM, Nikolenko SI, Pham S, Prjibelski AD, et al. 2012. SPAdes: a new genome assembly algorithm and its applications to single-cell sequencing. J. Comput. Biol. 19:455–477.

Bejerano G, Pheasant M, Makunin I, Stephen S, Kent WJ, Mattick JS, Haussler D. 2004. Ultraconserved elements in the human genome. Science 304:1321–1325.

Bi K, Vanderpool D, Singhal S, Linderoth T, Moritz C, Good JM. 2012. Transcriptome-based exon capture enables highly cost-effective comparative genomic data collection at moderate evolutionary scales. BMC Genom. 13:403.

Blom MPK, Bragg JG, Potter S, Moritz C. 2017. Accounting for uncertainty in gene tree estimation: summary-coalescent species tree inference in a challenging radiation of Australian lizards. Syst. Biol. 66:352–366.

Bragg JG, Potter S, Bi K, Moritz C. 2016. Exon capture phylogenomics: efficacy across scales of divergence. Mol. Ecol. Resour. 16:1059–1068.

Brandley MC, Bragg JG, Singhal S, Chapple DG, Jennings CK, Lemmon AR, Lemmon EM, Thompson MB, Moritz C. 2015. Evaluating the performance of anchored hybrid enrichment at the tips of the tree of life: a phylogenetic analysis of Australian *Eugongylus* group scincid lizards. BMC Evol. Biol. 15:62.

Broad Institute. Picard Tools. Version 2.20.1. http://broadinstitute.github.io/picard/

Brown RM, Siler CD, Richards SJ, Diesmos AC, Cannatella DC. 2015. Multilocus phylogeny and a new classification for Southeast Asian and Melanesian forest frogs (family Ceratobatrachidae). Zool. J. Linn. Soc. 174:130–168.

Bushnell B, Rood J, Singer E. 2017. BBMerge – Accurate paired shotgun read merging via overlap. PLoS ONE 12:e0185056–15.

Capella-Gutierrez S, Silla-Martinez JM, Gabaldon T. 2009. trimAl: A tool for automated alignment trimming in large-scale phylogenetic analyses. Bioinformatics 25:1972–1973.

Castoe TA, de Koning APJ, Kim H-M, Gu W, Noonan BP, Naylor G, Jiang ZJ, Parkinson CL, Pollock DD. 2009. Evidence for an ancient adaptive episode of convergent molecular evolution. Proc. Natl. Acad. Sci. USA 106:8986–8991.

Chakrabarty P, Faircloth BC, Alda F, Ludt WB, Mcmahan CD, Near TJ, Dornburg A, Albert JS, Arroyave J, Stiassny MLJ, et al. 2017. Phylogenomic systematics of ostariophysan fishes: ultraconserved elements support the surprising non-monophyly of Characiformes. Syst. Biol. 66:881–895.

Chang Z, Li G, Liu J, Zhang Y, Ashby C, Liu D, Cramer CL, Huang X. 2015. Bridger: A new framework for de novo transcriptome assembly using RNA-seq data. Genome Biol. 16:30.

Charif D, Lobry JR. 2007. SeqinR 1.0-2: A contributed package to the R project for statistical computing devoted to biological sequences retrieval and analysis. In: Structural approaches to sequence evolution. Vol. 3. Berlin: Springer. pp. 207–232; 26 p.

Chen S, Huang T, Zhou Y, Han Y, Xu M, Gu J. 2017. AfterQC: automatic filtering, trimming, error removing and quality control for fastq data. BMC Bioinf. 18:80.

Chen S, Zhou Y, Chen Y, Gu J. 2018 Mar 29. fastp: an ultra-fast all-in-one FASTQ preprocessor. bioRxiv:1–12. http://biorxiv.org/lookup/doi/10.1101/274100

Choi M, Scholl UI, Ji W, Liu T, Tikhonova IR, Zumbo P, Nayir A, Bakkaloğlu A, Ozen S, Sanjad S, et al. 2009. Genetic diagnosis by whole exome capture and massively parallel DNA sequencing. Proc. Natl. Acad. Sci. U.S.A. 106:19096–19101.

Crawford NG, Faircloth BC, McCormack JE, Brumfield RT, Winker K, Glenn TC. 2012. More than 1000 ultraconserved elements provide evidence that turtles are the sister group of archosaurs. Biol. Lett. 8:783–786.

Dool SE, Puechmaille SJ, Foley NM, Allegrini B, Bastian A, Mutumi GL, Maluleke TG, Odendaal LJ, Teeling EC, Jacobs DS. 2016. Nuclear introns outperform mitochondrial DNA in inter-specific phylogenetic reconstruction: Lessons from horseshoe bats (Rhinolophidae: Chiroptera). Mol. Phylogenet. Evol. 97:196–212.

Edwards SV. 2009. Natural selection and phylogenetic analysis. Proc. Natl. Acad. Sci. USA 106:8799–8800.

Edwards SV, Potter S, Schmitt CJ, Bragg JG, Moritz C. 2016. Reticulation, divergence, and the phylogeography–phylogenetics continuum. Proc. Natl. Acad. Sci. USA 113:8025–8032.

Faircloth BC, McCormack JE, Crawford NG, Harvey MG, Brumfield RT, Glenn TC. 2012. Ultraconserved elements anchor thousands of genetic markers spanning multiple evolutionary timescales. Syst. Biol. 61:717–726.

Feng YJ, Blackburn DC, Liang D, Hillis DM, Wake DB, Cannatella DC, Zhang P. 2017. Phylogenomics reveals rapid, simultaneous diversification of three major clades of Gondwanan frogs at the Cretaceous–Paleogene boundary. Proc. Natl. Acad. Sci. USA 114:E5864–E5870.

Frost DR, Grant T, Faivovich J, Bain RH, Haas A, Haddad C, De Sa RO, Channing A, Wilkinson M, Donnellan SC, et al. 2006. The amphibian tree of life. Bull. Am. Mus. Nat. Hist. 297:8–370.

Glaw F, Vences M. 2006. Phylogeny and genus-level classification of mantellid frogs (Amphibia, Anura). Org. Divers. Evol. 6:236–253.

Glenn TC. 2011. Field guide to next-generation DNA sequencers. Mol. Ecol. Resour. 11:759– 769.

Gnirke A, Melnikov A, Maguire J, Rogov P, LeProust EM, Brockman W, Fennell T, Giannoukos G, Fisher S, Russ C, et al. 2009. Solution hybrid selection with ultra-long oligonucleotides for massively parallel targeted sequencing. Nat. Biotechnol. 27:182–189.

Guillory WX, Muell MR, Summers K, Brown JL. 2019. Phylogenomic reconstruction of the Neotropical poison frogs (Dendrobatidae) and their conservation. Diversity 11:1–14.

Guo Y, Long J, He J, Li C-I, Cai Q, Shu X-O, Zheng W, Li C. 2012. Exome sequencing generates high quality data in non-target regions. BMC Genom. 13:194.

Hahn C, Bachmann L, Chevreux B. 2013. Reconstructing mitochondrial genomes directly from genomic next-generation sequencing reads–a baiting and iterative mapping approach. Nucleic Acids Res. 41:1–9.

Hahn MW, Nakhleh L. 2015. Irrational exuberance for resolved species trees. Evolution 70:7–17.

Halligan DL, Eyre-Walker A, Andolfatto P, Keightley PD. 2004. Patterns of evolutionary constraints in intronic and intergenic DNA of *Drosophila*. Genome Res. 14:273–279.

Han K-L, Braun EL, Kimball RT, Reddy S, Bowie RCK, Braun MJ, Chojnowski JL, Hackett SJ, Harshman J, Huddleston CJ, et al. 2011. Are transposable element insertions homoplasy free?: An examination using the avian tree of life. Syst. Biol. 60:375–386.

Hancock-Hanser BL, Frey A, Leslie MS, Dutton PH, Archer FI, Morin PA. 2013. Targeted multiplex next-generation sequencing: advances in techniques of mitochondrial and nuclear DNA sequencing for population genomics. Mol. Ecol. Resour. 13:254–268.

Hedge J, Wilson DJ. 2014. Bacterial phylogenetic reconstruction from whole genomes is robust to recombination but demographic inference is not. mBio 5:e02158–14.

Hedtke SM, Morgan MJ, Cannatella DC, Hillis DM. 2013. Targeted enrichment: Maximizing orthologous gene comparisons across deep evolutionary time. Joger U, editor. PLoS ONE 8:e67908.

Heinicke MP, Lemmon AR, Lemmon EM, McGrath K, Hedges SB. 2018. Phylogenomic support for evolutionary relationships of New World direct-developing frogs (Anura: Terraranae). Mol. Phylogenet. Evol. 118:145–155.

Hellsten U, Harland RM, Gilchrist MJ, Hendrix D, Jurka J, Kapitonov V, Ovcharenko I, Putnam NH, Shu S, Taher L, et al. 2010. The genome of the western clawed frog *Xenopus tropicalis*. Science 328:633–636.

Hobolth A, Dutheil JY, Hawks J, Schierup MH, Mailund T. 2011. Incomplete lineage sorting patterns among human, chimpanzee, and orangutan suggest recent orangutan speciation and widespread selection. Genome Res. 21:349–356.

Hosner PA, Braun EL, Kimball RT. 2016. Rapid and recent diversification of curassows, guans, and chachalacas (Galliformes: Cracidae) out of Mesoamerica: Phylogeny inferred from mitochondrial, intron, and ultraconserved element sequences. Mol. Phylogenet. Evol. 102:320–330.

Jones MR, Good JM. 2016. Targeted capture in evolutionary and ecological genomics. Mol. Ecol. 25:185–202.

Karin BR, Gamble T, Jackman TR. 2019. Optimizing phylogenomics with rapidly evolving long exons: Comparison with anchored hybrid enrichment and ultraconserved elements. bioRxiv 9:672238. http://biorxiv.org/lookup/doi/10.1101/672238

Katoh K, Standley DM. 2013. MAFFT multiple sequence alignment software version 7: Improvements in performance and usability. Mol. Biol. Evol. 30:772–780.

Katzman S, Kern AD, Bejerano G, Fewell G, Fulton L, Wilson RK, Salama SR, Haussler D. 2007. Human genome ultraconserved elements are ultraselected. Science 317:915–915.

Kent WJ. 2002. BLAT—the BLAST-like alignment tool. Genome Res. 12:656–664.

Kircher M, Kelso J. 2010. High-throughput DNA sequencing – concepts and limitations. Bioessays 32:524–536.

Knowles LL. 2009. Estimating species trees: Methods of phylogenetic analysis when there is incongruence across genes. Syst. Biol. 58:463–467.

Knowles LL, Huang H, Sukumaran J, Smith SA. 2018. A matter of phylogenetic scale: Distinguishing incomplete lineage sorting from lateral gene transfer as the cause of gene tree discord in recent versus deep diversification histories. Am. J. Bot. 105:376–384.

Lanier HC, Knowles LL. 2012. Is recombination a problem for species-tree analyses? Syst. Biol. 61:691–701.

Laurence M, Hatzis C, Brash DE. 2014. Common contaminants in next-generation sequencing that hinder discovery of low-abundance microbes. PLoS ONE 9:e97876–8.

Lawrence M, Huber W, Pagès H, Aboyoun P, Carlson M, Gentleman R, Morgan MT, Carey VJ. 2013. Software for computing and annotating genomic ranges. Prlic A, editor. PLOS Comput Biol 9:e1003118.

Leaché AD, Oaks JR. 2017. The utility of single nucleotide polymorphism (SNP) data in phylogenetics. Annu. Rev. Ecol. Evol. Syst. 48:69–84.

Leinonen R, Sugawara H, Shumway M, on behalf of the International Nucleotide Sequence Database Collaboration. 2010. The sequence read archive. Nucleic Acids Res. 39:D19–D21.

Lemmon AR, Emme SA, Lemmon EM. 2012. Anchored hybrid enrichment for massively high-throughput phylogenomics. Syst. Biol. 61:727–744.

Lemmon EM, Lemmon AR. 2013. High-throughput genomic data in systematics and phylogenetics. Annu. Rev. Ecol. Evol. Syst. 44:99–121.

Li H, Durbin R. 2010. Fast and accurate long-read alignment with Burrows–Wheeler transform. Bioinformatics 26:589–595.

Li H, Handsaker B, Wysoker A, Fennell T, Ruan J, Homer N, Marth G, Abecasis G, Durbin R, 1000 Genome Project Data Processing Subgroup. 2009. The Sequence Alignment/Map format and SAMtools. Bioinformatics 25:2078–2079.

Li W, Godzik A. 2006. Cd-hit: A fast program for clustering and comparing large sets of protein or nucleotide sequences. Bioinformatics 22:1658–1659.

Li Y-L, Weng J-C, Hsiao C-C, Chou M-T, Tseng C-W, Hung J-H. 2015. PEAT: An intelligent and efficient paired-end sequencing adapter trimming algorithm. BMC Bioinf. 16:S2.

Liu L, Yu L, Pearl DK, Edwards SV. 2009. Estimating species phylogenies using coalescence times among sequences. Syst. Biol. 58:468–477.

Maddison WP. 1997. Gene trees in species trees. Syst. Biol. 46:523–536.

McCartney-Melstad E, Mount GG, Shaffer HB. 2016. Exon capture optimization in amphibians with large genomes. Mol. Ecol. Resour. 16:1084–1094.

McCormack JE, Faircloth BC, Crawford NG, Gowaty PA, Brumfield RT, Glenn TC. 2012. Ultraconserved elements are novel phylogenomic markers that resolve placental mammal phylogeny when combined with species-tree analysis. Genome Res. 22:746–754.

McCormack JE, Hird SM, Zellmer AJ, Carstens BC, Brumfield RT. 2013. Applications of next-generation sequencing to phylogeography and phylogenetics. Mol. Phylogenet. Evol. 66:526–538.

McKenna A, Hanna M, Banks E, Sivachenko A, Cibulskis K, Kernytsky A, Garimella K, Altshuler D, Gabriel S, Daly M, et al. 2010. The Genome Analysis Toolkit: A MapReduce framework for analyzing next-generation DNA sequencing data. Genome Res. 20:1297– 1303.

Miller MR, Dunham JP, Amores A, Cresko WA, Johnson EA. 2007. Rapid and cost-effective polymorphism identification and genotyping using restriction site associated DNA (RAD) markers. Genome Res. 17:240–248.

Minh BQ, Nguyen MAT, Haeseler Von A. 2013. Ultrafast approximation for phylogenetic bootstrap. Mol. Biol. Evol. 30:1188–1195.

Mirarab S, Bayzid MS, Boussau B, Warnow T. 2014. Statistical binning enables an accurate coalescent-based estimation of the avian tree. Science 346:1250463–1250463.

Morgan M, Pagés H, Obenchain V, Hayden N. Rsamtools: Binary alignment (BAM), FASTA, variant call (BCF), and tabix file import.

Nguyen L-T, Schmidt HA, Haeseler von A, Minh BQ. 2014. IQ-TREE: A fast and effective stochastic algorithm for estimating maximum-likelihood phylogenies. Mol. Biol. Evol. 32:268–274.

Nikolenko SI, Korobeynikov AI, Alekseyev MA. 2013. BayesHammer: Bayesian clustering for error correction in single-cell sequencing. BMC Genom. 14:S7.

Paradis E, Schliep K. 2019. ape 5.0: An environment for modern phylogenetics and evolutionary analyses in R. Bioinformatics 35:526–528.

Peloso PLV, Frost DR, Richards SJ, Rodrigues MT, Donnellan S, Matsui M, Raxworthy CJ, Biju SD, Lemmon EM, Lemmon AR, et al. 2016. The impact of anchored phylogenomics and taxon sampling on phylogenetic inference in narrow-mouthed frogs (Anura, Microhylidae). Cladistics 32:113–140.

Pie MR, Faircloth BC, Ribeiro LF, Bornschein MR, McCormack JE. 2018. Phylogenomics of montane frogs of the Brazilian Atlantic Forest is consistent with isolation in sky islands followed by climatic stability. Biol. J. Linn. Soc. 125:72–82.

Portik DM, Smith LL, Bi K. 2016. An evaluation of transcriptome-based exon capture for frog phylogenomics across multiple scales of divergence (Class: Amphibia, Order: Anura). Mol. Ecol. Resour. 16:1069–1083.

Prum RO, Berv JS, Dornburg A, Field DJ, Townsend JP, Lemmon EM, Lemmon AR. 2015. A comprehensive phylogeny of birds (Aves) using targeted next-generation DNA sequencing. Nature 526:569–573.

Pyron RA, Wiens JJ. 2011. A large-scale phylogeny of Amphibia including over 2800 species, and a revised classification of extant frogs, salamanders, and caecilians. Mol. Phylogenet. Evol. 61:543–583.

Reddy S, Kimball RT, Pandey A, Hosner PA, Braun MJ, Hackett SJ, Han K-L, Harshman J, Huddleston CJ, Kingston S, et al. 2017. Why do phylogenomic data sets yield conflicting trees? Data type influences the avian tree of life more than taxon sampling. Syst. Biol. 66:857–879.

Reilly SB, Stubbs AL, Karin BR, Bi K, Arida E, Iskandar DT, McGuire JA. 2019. Leap-frog dispersal and mitochondrial introgression: Phylogenomics and biogeography of Limnonectes fanged frogs in the Lesser Sundas Archipelago of Wallacea. J. Biogeogr. 46:757–769.

Richards EJ, Brown JM, Barley AJ, Chong RA, Thomson RC. 2018. Variation across mitochondrial gene trees provides evidence for systematic error: How much gene tree variation Is biological? Syst. Biol. 67:847–860.

Rognes T, Flouri T, Nichols B, Quince C, Mahé F. 2016. VSEARCH: A versatile open source tool for metagenomics. PeerJ 4:e2584–22.

Rohland N, Reich D. 2012. Cost-effective, high-throughput DNA sequencing libraries for multiplexed target capture. Genome Res. 22:939–946.

Safonova Y, Bankevich A, Pevzner PA. 2015. dipSPAdes: Assembler for highly polymorphic diploid genomes. J. Comput. Biol. 22:528–545.

Shendure J, Ji H. 2008. Next-generation DNA sequencing. Nat. Biotechnol. 26:1135–1145.

Simmons MP, Ochoterena H. 2000. Gaps as characters in sequence-based phylogenetic analyses. Syst. Biol. 49:369–381.

Singhal S, Grundler M, Colli G, Rabosky DL. 2017. Squamate Conserved Loci (SqCL): A unified set of conserved loci for phylogenomics and population genetics of squamate reptiles. Mol. Ecol. Resour. 17:e12–e24.

Smith BT, Harvey MG, Faircloth BC, Glenn TC, Brumfield RT. 2014. Target capture and massively parallel sequencing of ultraconserved elements for comparative studies at shallow evolutionary time scales. Syst. Biol. 63:83–95.

Springer MS, Gatesy J. 2018. Delimiting coalescence genes (C-Genes) in phylogenomic data sets. Genes 9:1–19.

Stephen S, Pheasant M, Makunin IV, Mattick JS. 2008. Large-scale appearance of ultraconserved elements in tetrapod genomes and slowdown of the molecular Clock. Mol. Biol. Evol. 25:402–408.

Streicher JW, Miller EC, Guerrero PC, Correa C, Ortiz JC, Crawford AJ, Pie MR, Wiens JJ. 2018. Evaluating methods for phylogenomic analyses, and a new phylogeny for a major frog clade (Hyloidea) based on 2214 loci. Mol. Phylogenet. Evol. 119:128–143.

Sulonen A-M, Ellonen P, Almusa H, Lepistö M, Eldfors S, Hannula S, Miettinen T, Tyynismaa H, Salo P, Heckman C, et al. 2011. Comparison of solution-based exome capture methods for next generation sequencing. Genome Biol. 12:R94.

Sun Y-B, Xiong Z-J, Xiang X-Y, Liu S-P, Zhou W-W, Tu X-L, Zhong L, Wang L, Wu D-D, Zhang B-L, et al. 2015. Whole-genome sequence of the Tibetan frog *Nanorana parkeri* and the comparative evolution of tetrapod genomes. Proc. Natl. Acad. Sci. USA 112:E1257– E1262.

Taylor DJ, Piel WH. 2004. An assessment of accuracy, error, and conflict with support values from genome-scale phylogenetic data. Mol. Biol. Evol. 21:1534–1537.

Tewhey R, Nakano M, Wang X, Pabón-Peña C, Novak B, Giuffre A, Lin E, Happe S, Roberts DN, LeProust EM, et al. 2009. Enrichment of sequencing targets from the human genome by solution hybridization. Genome Biol. 10:R116.

Townsend J. 2007. Profiling phylogenetic informativeness. Syst. Biol. 56:222–231.

Van der Auwera GA, Carneiro MO, Hartl C, Poplin R, del Angel G, Moonshine AL, Jordan T, Shakir K, Roazen D, Thibault J, et al. 2013. From FastQ data to high-confidence variant calls: The genome analysis toolkit best practices pipeline. Curr. Prot. Bioinf. 11:11.10.1–11.10.33.

Wang Z, Gerstein M, Snyder M. 2009. RNA-Seq: A revolutionary tool for transcriptomics. Nat. Rev. Genet. 10:57–63.

Xi Z, Liu L, Davis CC. 2015. Genes with minimal phylogenetic information are problematic for coalescent analyses when gene tree estimation is biased. Mol. Phylogenet. Evol. 92:63–71.

Yuan Z-Y, Zhang B-L, Raxworthy CJ, Weisrock DW, Hime PM, Jin J-Q, Lemmon EM, Lemmon AR, Holland SD, Kortyna ML, et al. 2019. Natatanuran frogs used the Indian Plate to step-stone disperse and radiate across the Indian Ocean. Nat. Sci. Rev. 6:10–14.

Zarza E, Faircloth BC, Tsai WLE, Bryson RW Jr, Klicka J, McCormack JE. 2016. Hidden histories of gene flow in highland birds revealed with genomic markers. Mol. Ecol. 25:5144–5157.

