## Supplementary Materials for "FrogCap: A modular sequence capture probe set for phylogenomics and population genetics for all frogs, assessed across multiple phylogenetic scales"

From:

**Table S1.** The transcriptomes and their associated GenBank ID used in this study are shown. Assembled transcriptomes will be available in the Open Science Framework (https://osf.io/gvbr5/) following manuscript acceptance.

| SuperFamily | Family | Species | GenBank Accession Number |
|---|---|---|---|
| Hyloidea | Hylidae | *Agalychnis callidryas* | PRJNA259743 |
| Hyloidea | Bufonidae | *Atelopus glyphus* | PRJNA259720 |
| Hyloidea | Bufonidae | *Atelopus zeteki* | PRJNA216050 |
| Basal | Bombinatoridae | *Bombina maxima* | PRJNA174732 |
| Hyloidea | Bufonidae | *Bufo marinus* | PRJNA395127 |
| Hyloidea | Bufonidae | *Bufo schneideri* | PRJNA255079 |
| Hyloidea | Bufonidae | *Bufo spinulosus* | PRJNA236569 |
| Hyloidea | Craugastoridae | *Craugastor fitzingeri* | PRJNA259742 |
| Hyloidea | Hylidae | *Cyclorana alboguttata* | PRJNA177363 |
| Hyloidea | Centrolenidae | *Espadarana prosoblepon* | PRJNA237648 |
| Hyloidea | Hylidae | *Hyla arborea* | PRJNA196399 |
| Ranoidea | Hyperoliidae | *Kassina decorata* | Unpublished |
| Ranoidea | Arthroleptidae | *Leptodactylodon ovatus* | Unpublished |
| Ranoidea | Hyperoliidae | *Afrixalus paradorsalis* | Unpublished |
| Ranoidea | Arthroleptidae | *Leptopelis boulengeri* | Unpublished |
| Ranoidea | Dicroglossidae | *Nanorana parkeri* | PRJNA344660 |
| Ranoidea | Ranidae | *Pelophylax lessonae* | PRJNA230915 |
| Ranoidea | Ranidae | *Pelophylax nigromaculatus* | PRJNA299518 |
| Ranoidea | Rhacophoridae | *Polypedates megacephalus* | PRJNA299518 |
| Ranoidea | Hylidae | *Pseudacris regilla* | PRJNA163143 |
| Ranoidea | Dicroglossidae | *Quasipaa boulengeri* | PRJNA304335 |
| Ranoidea | Ranidae | *Rana chensinensis* | PRJNA178186 |
| Ranoidea | Ranidae | *Rana clamitans* | PRJNA162931 |
| Ranoidea | Ranidae | *Rana clamitans* | PRJNA240240 |
| Ranoidea | Ranidae | *Rana kukunoris* | PRJNA178186 |

| | | | |
|---|---|---|---|
| Ranoidea | Ranidae | *Rana margaretae* | PRJNA299518 |
| Ranoidea | Ranidae | *Rana sylvatica* | PRJNA392411 |
| Ranoidea | Ranidae | *Rana temporaria* | PRJEB9622 |
| Ranoidea | Ranidae | *Rana yavapaiensis* | PRJNA232044 |
| Ranoidea | Rhacophoridae | *Rhacophorus dennysi* | PRJNA299518 |

**Table S2.** The samples and their associated collection and museum ID used in this study are shown. Samples included in multiple phylogenetic scales are only listed once.

| Probe Set | Scale | Species | Collection ID | Museum ID | GenBank Accession |
|---|---|---|---|---|---|
| Ranoidea | Species | *Cornufer vertebralis* | A 24L | - | Manuscript acceptance |
| Ranoidea | Species | *Cornufer vertebralis* | - | AMNH A-168972 | Data for outgroups |
| Ranoidea | Species | *Cornufer vertebralis* | JLW 173 | KU 349559 | Available upon request |
| Ranoidea | Species/ Genus | *Cornufer vertebralis* | JLW 343 | KU 349545 | |
| Ranoidea | Species | *Cornufer vertebralis* | JQR 1793 | KU 341540 | |
| Ranoidea | Species | *Cornufer vertebralis* | JQR 1868 | KU 341190 | |
| Ranoidea | Species | *Cornufer vertebralis* | JQR 1869 | MCZ A-149365 | |
| Ranoidea | Species | *Cornufer vertebralis* | JQR2001 | MCZ A-151173 | |
| Ranoidea | Species | *Cornufer vertebralis* | RMB 7128 | KU 307406 | |
| Ranoidea | Species | *Cornufer vertebralis* | SLT 179 | KU 341198 | |
| Ranoidea | Species | *Cornufer vertebralis* | SLT 200 | KU 341191 | |
| Ranoidea | Species | *Cornufer vertebralis* | SLT 203 | MCZ A-149366 | |
| Ranoidea | Species | *Cornufer vertebralis* | SLT 268 | MCZ A-149388 | |
| Ranoidea | Species | *Cornufer vertebralis* | SLT 280 | KU 341192 | |
| Ranoidea | Species | *Cornufer vertebralis* | SLT 524 | KU 343643 | |
| Ranoidea | Species | *Cornufer vertebralis* | SLT 674 | KU 343653 | |
| Ranoidea | Superfamily/ Family | *Aglyptodactylus securifer* | CRH1644 | KU 347232 | |
| Ranoidea | Superfamily | *Amietia angolensis* | - | CAS 254876 | |
| Ranoidea | Superfamily | *Amolops ricketti* | KUFS65 | KU 311638 | |
| Ranoidea | Superfamily | *Anodontohyla boulengeri* | CRH1898 | KU 347449 | |
| Ranoidea | Superfamily | *Arthroleptis variabilis* | RMB19372 | KU 341900 | |
| Ranoidea | Superfamily/ Family | *Boophis tephraeomystax* | CRH1675 | KU 347320 | |
| Ranoidea | Superfamily | *Cardioglossa leucomystax* | RMB19294 | KU 341857 | |

| | | | | |
|---|---|---|---|---|
| Ranoidea | Superfamily/ Genus | *Cornufer guentheri* | SLT345 | KU 341235 |
| Ranoidea | Superfamily/ Genus | *Cornufer gilliardi* | JF112 | - |
| Ranoidea | Superfamily | *Gastrophyrne olivacae* | KUFS914 | KU 342724 |
| Ranoidea | Superfamily | *Heterixalus punctatus* | CRH1685 | KU 347213 |
| Ranoidea | Superfamily | *Hylarana erythraea* | RMB4300 | PNM |
| Ranoidea | Superfamily | *Ingerana mariae* | RMB7803 | KU 309518 |
| Ranoidea | Superfamily | *Kassina senegalensis* | RAX 2025 | KU 290414 |
| Ranoidea | Superfamily | *Leptodactylodon ovatus* | - | CAS 253933 |
| Ranoidea | Superfamily | *Limnonectes kuhlii* | RMB2127 | PNM |
| Ranoidea | Superfamily | *Microhyla heymonsi* | KUFS265 | KU 311616 |
| Ranoidea | Superfamily | *Nyctixalus pictus* | - | MVZ 239460 |
| Ranoidea | Superfamily | *Odontobatrachus natator* | CAS230052 | CAS 230052 |
| Ranoidea | Superfamily | *Oreophryne annulata* | RMB10029 | KU 314333 |
| Ranoidea | Superfamily | *Platymantis corrugatus* | RMB15045 | KU 330258 |
| Ranoidea | Superfamily | *Pulchrana moellendorffi* | ACD1277 | KU 327049 |
| Ranoidea | Superfamily | *Rhacophorus bipunctatus* | - | MVZ 221351 |
| Ranoidea | Superfamily | *Scaphiophryne marmorata* | CRH920 | KU 343458 |
| Ranoidea | Family | *Blommersia grandisonae* | CRH792 | KU 340891 |
| Ranoidea | Family | *Gephyromantis redimus* | CRH1628 | KU 347352 |
| Ranoidea | Family | *Guibemantis depressiceps* | CRH535 | KU 340774 |
| Ranoidea | Family | *Mantella baroni* | CRH1027 | KU 343297 |
| Ranoidea | Family | *Mantidactylus melanopleura* | CRH1998 | KU 347429 |
| Ranoidea | Family | *Spinomantis bertini* | CRH726 | KU 340852 |
| Ranoidea | Genus | *Cornufer batantae* | SJR7502 | - |
| Ranoidea | Genus | *Cornufer boulengeri* | FK10744 | BPBM 22329 |
| Ranoidea | Genus | *Cornufer bufoniformis* | SJR5475 | - |
| Ranoidea | Genus | *Cornufer cryptotis yapeni* | RG7979 | ZMB 80356 |

| | | | | |
|---|---|---|---|---|
| Ranoidea | Genus | *Cornufer custos* | ABTC136625 | - |
| Ranoidea | Genus | *Cornufer exedrus* | SJR10784 | SAMA R64760 |
| Ranoidea | Genus | *Cornufer guppyi* | SLT679 | KU 343719 |
| Ranoidea | Genus | *Cornufer hedigeri* | SLT710 | KU 343799 |
| Ranoidea | Genus | *Cornufer heffernani* | SJR5303 | SAMA R56799 |
| Ranoidea | Genus | *Cornufer latro* | SJR2145 | SAMA R62821 |
| Ranoidea | Genus | *Cornufer magnus* | SJR5220 | SAMA R60238 |
| Ranoidea | Genus | *Cornufer minutus* | SLT202 | KU 341495 |
| Ranoidea | Genus | *Cornufer papuensis* | RG7373 | - |
| Ranoidea | Genus | *Cornufer parkeri* | CCA2644 | LSUMZ 93781 |
| Ranoidea | Genus | *Cornufer punctatus* | RG7970 | - |
| Ranoidea | Genus | *Cornufer schmidti* | CCA1583 | LSUMZ 94179 |
| Ranoidea | Genus | *Cornufer solomonis* | RMB6960 | KU 307238 |
| Ranoidea | Genus | *Cornufer trossulus* | SLT262 | KU 341538 |
| Ranoidea | Genus | *Cornufer vitianus* | CM1498 | - |
| Ranoidea | Genus | *Cornufer weberi* | SLT269 | KU 341569 |
| Ranoidea | Genus | *Cornufer wuenscheorum* | RG7751 | ZMB 67214 |
| Reduced | Reduced | *Occidozyga cf. laevis* | CDS2282 | KU 308045 |
| Reduced | Reduced | *Occidozyga cf. laevis* | CDS1257 | KU 302322 |
| Reduced | Reduced | *Occidozyga cf. laevis* | CDS1808 | KU 306299 |
| Reduced | Reduced | *Occidozyga cf. laevis* | CDS2026 | KU 306277 |
| Reduced | Reduced | *Occidozyga cf. laevis* | CDS2045 | KU 306232 |
| Reduced | Reduced | *Occidozyga cf. laevis* | CDS270 | KU 202245 |
| Reduced | Reduced | *Occidozyga cf. laevis* | CDS3582 | KU 320163 |
| Reduced | Reduced | *Occidozyga cf. laevis* | CDS487 | KU 302291 |
| Reduced | Reduced | *Occidozyga cf. laevis* | CDS543 | KU 302274 |
| Reduced | Reduced | *Occidozyga cf. laevis* | RMB11020 | KU 321227 |
| Reduced | Reduced | *Occidozyga cf. laevis* | RMB15467 | KU 333853 |
| Reduced | Reduced | *Occidozyga cf. laevis* | RMB16321 | KU 333098 |

| | | | | |
|---|---|---|---|---|
| Reduced | Reduced | *Occidozyga cf. laevis* | RMB16436 | KU 333103 |
| Reduced | Reduced | *Occidozyga cf. laevis* | RMB17235 | KU 334639 |
| Reduced | Reduced | *Occidozyga cf. laevis* | RMB20282 | KU 345760 |
| Reduced | Reduced | *Occidozyga cf. laevis* | RMB20559 | KU 345738 |
| Reduced | Reduced | *Occidozyga cf. laevis* | RMB2133 | PNM |
| Reduced | Reduced | *Occidozyga cf. laevis* | RMB2542 | PNM |
| Reduced | Reduced | *Occidozyga cf. laevis* | RMB3068 | KU 326484 |
| Reduced | Reduced | *Occidozyga cf. laevis* | RMB3127 | PNM |
| Reduced | Reduced | *Occidozyga cf. laevis* | RMB4318 | PNM |
| Reduced | Reduced | *Occidozyga cf. laevis* | RMB4468 | THNC 63050 |
| Reduced | Reduced | *Occidozyga cf. laevis* | RMB5396 | KU 303492 |
| Reduced | Reduced | *Occidozyga cf. laevis* | RMB6117 | KU 307540 |
| Reduced | Reduced | *Occidozyga cf. laevis* | RMB6411 | KU 306917 |
| Reduced | Reduced | *Occidozyga cf. laevis* | RMB7686 | KU 309480 |
| Reduced | Reduced | *Occidozyga cf. laevis* | RMB819 | PNM |
| Reduced | Reduced | *Occidozyga cf. laevis* | RMB8750 | KU 315273 |
| Reduced | Reduced | *Occidozyga cf. laevis* | RMB8900 | KU 326227 |
| Reduced | Reduced | *Occidozyga cf. laevis* | RMB9600 | KU 313686 |
| Hyloidea | Superfamily | *Allobates femoralis* | WED55592 | KU 205297 |
| Hyloidea | Superfamily | *Anaryxus woodhousii* | RMB19577 | KU 341629 |
| Hyloidea | Superfamily | *Ansonia mcgregori* | RMB10371 | KU 314180 |
| Hyloidea | Superfamily | *Barycholos pulcher* | LAC692 | KU 217780 |
| Hyloidea | Superfamily | *Bolitaglossa palmata* | LAC1154 | KU 217423 |
| Hyloidea | Superfamily | *Incilius nebulifer* | EAL 5027 | KU 203846 |
| Hyloidea | Superfamily | *Ceratophrys cornuta* | WED57689 | KU 215537 |
| Hyloidea | Superfamily | *Craugastor rupinius* | JES2401 | KU 291264 |
| Hyloidea | Superfamily | *Dendrobates histrionicus* | LAC2539 | KU 223498 |
| Hyloidea | Superfamily | *Eluetherodactylus sp* | GLOR6202 | KU-Uncatalogued |
| Hyloidea | Superfamily | *Gastrotheca marsupiata* | WED58521 | KU 214816 |

| | | | | |
|---|---|---|---|---|
| Hyloidea | Superfamily | *Hyalinobatrachium fleischmanni* | - | MZUTI 3621 |
| Hyloidea | Superfamily | *Hyloscirtus phyllognathus* | WED58378 | KU 212118 |
| Hyloidea | Superfamily | *Leptobrachium hainensis* | KUFS192 | KU 311582 |
| Hyloidea | Superfamily | *Leptodactylus pentadactylus* | WED55494 | KU 205193 |
| Hyloidea | Superfamily | *Nyctimystes infrafrenatus* | SLT771 | KU 345125 |
| Hyloidea | Superfamily | *Oreobates quizensis* | WED59885 | KU 222033 |
| Hyloidea | Superfamily | *Phyrnohyas venulosa* | WED55450 | KU 205418 |
| Hyloidea | Superfamily | *Physalamus pustulatus* | LAC1642 | KU 218219 |
| Hyloidea | Superfamily | *Pristimantis w-nigrum* | WED53045 | KU 202561 |
| Hyloidea | Superfamily | *Spea bombifrons* | RMB19586 | KU 341688 |
| Hyloidea | Superfamily | *Strabomantis sulcatus* | LAC841 | KU 218056 |
| Hyloidea | Superfamily | *Telmatobius truebae* | WED56933 | KU 212465 |
| Hyloidea | Superfamily | *Vitreorana castroviejoi* | JMG33 | MHNLS 16446 |

**Table S3.** The genomes for the raw read decontamination step and their associated GenBank ID used in this study are shown.

| Group | Genome | GenBank_Accession |
|---|---|---|
| Database | UniVec | https://www.ncbi.nlm.nih.gov/tools/vecscreen/univec/ |
| Animal | Caenorhabditis | GCA_000002985.3 |
| Animal | Drosophila | GCA_000001215.4 |
| Bacteria | Achromobacter | GCA_001971645.1 |
| Bacteria | Acidaminococcus | GCA_000025305.1 |
| Bacteria | Acinetobacter | GCA_002055515.1 |
| Bacteria | Afipia | GCA_000314735.2 |
| Bacteria | Agrobacterium | GCA_000016265.1 |
| Bacteria | Alcaligenes | GCA_000967305.2 |
| Bacteria | Aminobacter | GCA_001605015.1 |
| Bacteria | Bradyrhizobium | GCA_900011245.1 |
| Bacteria | Brevundimonas | GCA_000635915.2 |
| Bacteria | Burkholderia | GCA_001999785.1 |
| Bacteria | Corynebacterium | GCA_002804085.1 |
| Bacteria | Curvibacter | GCA_000381265.1 |
| Bacteria | Escherichia | GCA_000008865.2 |
| Bacteria | Flavobacterium | GCA_000064305.2 |
| Bacteria | Haemophilus | GCA_003390455.1 |
| Bacteria | Helcococcus | GCA_000245755.1 |
| Bacteria | Herbaspirillum | GCA_001267925.1 |
| Bacteria | Legionella | GCA_000008485.1 |
| Bacteria | Leifsonia | GCA_000470775.1 |
| Bacteria | Mesorhizobium | GCA_000185905.1 |
| Bacteria | Methylobacterium | GCA_000022085.1 |
| Bacteria | Microbacterium | GCA_000202635.1 |
| Bacteria | Moraxella | GCA_001553955.1 |

| | | |
|---|---|---|
| Bacteria | Mycoplasma | GCA_000011445.1 |
| Bacteria | Novosphingobium | GCA_000767465.1 |
| Bacteria | Ochrobactrum | GCA_000017405.1 |
| Bacteria | Pedobacter | GCA_000023825.1 |
| Bacteria | Phyllobacterium | GCA_002764115.1 |
| Bacteria | Pseudomonas | GCA_000007805.1 |
| Bacteria | Pseudonocardia | GCA_000196675.1 |
| Bacteria | Ralstonia | GCA_000020205.1 |
| Bacteria | Rhodococcus | GCA_000982715.1 |
| Bacteria | Salmonella | GCA_000006945.2 |
| Bacteria | Sphingomonas | GCA_000512205.2 |
| Bacteria | Staphylococcus | GCA_000013425.1 |
| Bacteria | Stenotrophomonas | GCA_000072485.1 |
| Bacteria | Streptococcus | GCA_000014205.1 |
| Fungus | Aspergillus | GCA_000149205.2 |
| Fungus | Magnaporthe | GCA_001936955.1 |
| Fungus | Malassezia | GCA_003290485.1 |
| Fungus | Penicillium | GCA_000710275.1 |
| Fungus | Puccinia | GCA_001624995.1 |
| Fungus | Saccharomyces | GCA_000146045.2 |
| Fungus | Schizosaccharomyces | GCA_000002945.2 |

**Table S4.** The samples and their associated collection ID used in this study are shown. Each samples' summary statistics are shown, where orthologs and paralogs are discovered during probe matching. The length statistics are calculated from the orthologous contigs.

| Probe Set | Scale | Sample | N contigs | N orthologs | N paralogs | Min length | Max length | Mean length | Median length |
|---|---|---|---|---|---|---|---|---|---|
| Ranoidea | Species | *Cornufer vertebralis* A 24L | 17583 | 11181 | 442 | 125 | 5840 | 845.7 | 795 |
| Ranoidea | Species | *Cornufer vertebralis* A168972 | 12690 | 11096 | 542 | 129 | 5653 | 921.9 | 894 |
| Ranoidea | Species | *Cornufer vertebralis* JLW 173 | 14253 | 11088 | 594 | 104 | 4892 | 827.9 | 787.5 |
| Ranoidea | Species | *Cornufer vertebralis* JLW 343 | 16767 | 11557 | 692 | 167 | 7547 | 912.0 | 874 |
| Ranoidea | Species | *Cornufer vertebralis* JQR 1793 | 14248 | 11018 | 640 | 84 | 5728 | 811.4 | 767 |
| Ranoidea | Species | *Cornufer vertebralis* JQR 1868 | 16580 | 10736 | 509 | 93 | 7339 | 773.6 | 730 |
| Ranoidea | Species | *Cornufer vertebralis* JQR 1869 | 11098 | 10010 | 314 | 116 | 9788 | 803.7 | 761 |
| Ranoidea | Species | *Cornufer vertebralis* JQR2001 | 13788 | 11026 | 585 | 101 | 4236 | 835.8 | 791 |
| Ranoidea | Species | *Cornufer vertebralis* RMB 7128 | 16344 | 9666 | 271 | 118 | 7266 | 784.1 | 724.5 |
| Ranoidea | Species | *Cornufer vertebralis* SLT 179 | 12983 | 11010 | 496 | 109 | 12413 | 850.4 | 808 |
| Ranoidea | Species | *Cornufer vertebralis* SLT 200 | 12080 | 10798 | 393 | 117 | 12118 | 810.8 | 771 |
| Ranoidea | Species | *Cornufer vertebralis* SLT 203 | 16007 | 9692 | 299 | 93 | 4517 | 767.2 | 716 |
| Ranoidea | Species | *Cornufer vertebralis* SLT 268 | 12655 | 10066 | 325 | 129 | 12173 | 829.8 | 782 |
| Ranoidea | Species | *Cornufer vertebralis* SLT 280 | 14160 | 11106 | 640 | 107 | 7124 | 845.1 | 807 |
| Ranoidea | Species | *Cornufer vertebralis* SLT 524 | 11810 | 10538 | 350 | 61 | 12309 | 825.9 | 787 |
| Ranoidea | Species | *Cornufer vertebralis* SLT 674 | 14742 | 10823 | 539 | 120 | 12713 | 875.7 | 832 |
| Ranoidea | Superfamily | *Aglyptodactylus securifer* CRH1644 | 12005 | 11118 | 453 | 76 | 12498 | 965.6 | 936 |
| Ranoidea | Superfamily | *Amietia angolensis* CAS254876 | 13943 | 11989 | 647 | 138 | 9433 | 944.8 | 919 |
| Ranoidea | Superfamily | *Amolops ricketti* KUFS65 | 9527 | 8628 | 329 | 92 | 12583 | 767.1 | 699 |
| Ranoidea | Superfamily | *Anodontohyla boulengeri* CRH1898 | 9516 | 7111 | 284 | 106 | 23144 | 815.2 | 765 |
| Ranoidea | Superfamily | *Arthroleptis variabilis* RMB19372 | 8530 | 6147 | 267 | 111 | 6045 | 821.7 | 776 |
| Ranoidea | Superfamily | *Boophis tephraeomystax* CRH1675 | 14168 | 11198 | 405 | 129 | 15946 | 1114.2 | 1082 |
| Ranoidea | Superfamily | *Cardioglossa leucomystax* RMB19294 | 9117 | 6430 | 248 | 98 | 12048 | 819.9 | 775 |
| Ranoidea | Superfamily | *Cornufer guentheri* SLT345 | 13498 | 11695 | 470 | 147 | 6840 | 930.6 | 899 |
| Ranoidea | Superfamily | *Cornufer gilliardi* JF112 | 15155 | 11752 | 723 | 79 | 12400 | 957.9 | 921 |

| | | | | | | | | | |
|---|---|---|---|---|---|---|---|---|---|
| Ranoidea | Superfamily | *Gastrophyrne olivacae* KUFS914 | 8949 | 6026 | 172 | 112 | 11909 | 848.8 | 802 |
| Ranoidea | Superfamily | *Heterixalus punctatus* CRH1685 | 15189 | 8681 | 356 | 110 | 6692 | 908.2 | 869 |
| Ranoidea | Superfamily | *Hylarana erythraea* RMB4300 | 12230 | 10783 | 640 | 101 | 12268 | 929.5 | 902 |
| Ranoidea | Superfamily | *Ingerana mariae* RMB7803 | 12329 | 11311 | 473 | 145 | 18047 | 958.0 | 928 |
| Ranoidea | Superfamily | *Kassina senegalensis* KU290414 | 8562 | 6986 | 257 | 110 | 12075 | 811.2 | 772 |
| Ranoidea | Superfamily | *Leptodactylodon ovatus* CAS253933 | 9873 | 8276 | 284 | 81 | 5529 | 875.5 | 830 |
| Ranoidea | Superfamily | *Limnonectes kuhlii* RMB2127 | 15648 | 12235 | 865 | 156 | 10844 | 1005.6 | 981 |
| Ranoidea | Superfamily | *Microhyla heymonsi* KUFS265 | 8023 | 5225 | 198 | 80 | 11467 | 838.2 | 784 |
| Ranoidea | Superfamily | *Nyctixalus pictus* 239460 | 14308 | 11710 | 560 | 115 | 8314 | 1003.5 | 962 |
| Ranoidea | Superfamily | *Odontobatrachus natator* CAS230052 | 12659 | 11865 | 403 | 122 | 12170 | 892.2 | 862 |
| Ranoidea | Superfamily | *Oreophryne annulata* RMB10029 | 6968 | 5050 | 85 | 102 | 12605 | 791.3 | 742 |
| Ranoidea | Superfamily | *Platymantis corrugatus* RMB15045 | 11706 | 10571 | 333 | 106 | 10810 | 858.5 | 826 |
| Ranoidea | Superfamily | *Pulchrana moellendorffi* KU 327049 | 18062 | 11770 | 911 | 103 | 12633 | 998.9 | 970 |
| Ranoidea | Superfamily | *Rhacophorus bipunctatus* 221351 | 16015 | 11745 | 777 | 114 | 9515 | 982.9 | 947 |
| Ranoidea | Superfamily | *Scaphiophryne marmorata* CRH920 | 14222 | 9007 | 651 | 111 | 22883 | 868.4 | 831 |
| Ranoidea | Superfamily | *Aglyptodactylus securifer* CRH1644 | 12005 | 11118 | 453 | 76 | 12498 | 965.6 | 936 |
| Ranoidea | Superfamily | *Blommersia grandisonae* CRH792 | 12170 | 10697 | 515 | 125 | 5288 | 895.8 | 858 |
| Ranoidea | Superfamily | *Boophis tephraeomystax* CRH1675 | 14168 | 11198 | 405 | 129 | 15946 | 1114.2 | 1082 |
| Ranoidea | Superfamily | *Gephyromantis redimitus* CRH1628 | 14178 | 11259 | 1042 | 96 | 10021 | 901.1 | 872 |
| Ranoidea | Superfamily | *Guibemantis depressiceps* CRH535 | 12747 | 11017 | 503 | 119 | 9668 | 1044.5 | 1011 |
| Ranoidea | Superfamily | *Mantella baroni* CRH1027 | 12099 | 9428 | 347 | 122 | 12213 | 964.5 | 919 |
| Ranoidea | Superfamily | *Mantidactylus melanopleura* CRH1998 | 12865 | 10928 | 899 | 101 | 16850 | 917.5 | 882 |
| Ranoidea | Superfamily | *Spinomantis bertini* CRH726 | 13326 | 11920 | 630 | 129 | 12744 | 1107.9 | 1084 |
| Ranoidea | Genus | *Cornufer batantae* SJR 7502 | 12575 | 9783 | 292 | 101 | 12727 | 881.6 | 824 |
| Ranoidea | Genus | *Cornufer boulengeri* FK 10744 | 14132 | 10440 | 326 | 94 | 12322 | 889.0 | 826 |
| Ranoidea | Genus | *Cornufer bufoniformis* SJR 5475 | 13601 | 11066 | 418 | 97 | 12190 | 1006.2 | 971 |
| Ranoidea | Genus | *Cornufer cryptotis yapeni* RG 7979 | 13980 | 10970 | 396 | 146 | 7935 | 1032.1 | 987 |
| Ranoidea | Genus | *Cornufer custos* ABTC 136625 | 19919 | 11064 | 447 | 142 | 12217 | 1017.9 | 964 |
| Ranoidea | Genus | *Cornufer exedrus* SJR 10784 | 10062 | 8356 | 223 | 61 | 12265 | 782.1 | 719 |

| | | | | | | | | | |
|---|---|---|---|---|---|---|---|---|---|
| Ranoidea | Genus | *Cornufer gilliardi* JF 112 | 15111 | 11913 | 622 | 144 | 12400 | 974.3 | 942 |
| Ranoidea | Genus | *Cornufer guentheri* SLT 345 | 13770 | 11569 | 603 | 143 | 14988 | 889.3 | 861 |
| Ranoidea | Genus | *Cornufer guppyi* SLT 679 | 12322 | 11320 | 596 | 142 | 4943 | 1001.6 | 976 |
| Ranoidea | Genus | *Cornufer hedigeri* SLT 710 | 12670 | 11542 | 511 | 62 | 8940 | 947.1 | 916 |
| Ranoidea | Genus | *Cornufer heffernani* 5303 5316 | 11591 | 9772 | 319 | 122 | 12134 | 881.2 | 823 |
| Ranoidea | Genus | *Cornufer latro* SR 2145 | 13005 | 10862 | 445 | 124 | 6006 | 828.4 | 790 |
| Ranoidea | Genus | *Cornufer magnus* 5220 5252 | 12439 | 11018 | 412 | 129 | 12326 | 879.8 | 838 |
| Ranoidea | Genus | *Cornufer minutus* SLT 202 | 12495 | 10831 | 457 | 119 | 8937 | 801.6 | 763 |
| Ranoidea | Genus | *Cornufer papuensis* RG 7373 | 11936 | 10954 | 491 | 116 | 5854 | 884.6 | 855 |
| Ranoidea | Genus | *Cornufer parkeri* CCA 2644 | 13333 | 10288 | 523 | 69 | 5899 | 814.5 | 764 |
| Ranoidea | Genus | *Cornufer punctatus* RG 7970 | 12013 | 10973 | 386 | 166 | 12435 | 933.2 | 897 |
| Ranoidea | Genus | *Cornufer schmidti* CCA 1583 | 11540 | 10500 | 350 | 115 | 5928 | 887.0 | 854 |
| Ranoidea | Genus | *Cornufer solomonis* RMB 6960 | 35201 | 11388 | 538 | 186 | 15024 | 1206.4 | 1150 |
| Ranoidea | Genus | *Cornufer trossulus* SLT 262 | 12253 | 10168 | 295 | 144 | 14170 | 805.7 | 749 |
| Ranoidea | Genus | *Cornufer vertebralis* JLW 343 | 16767 | 11557 | 692 | 167 | 7547 | 912.0 | 874 |
| Ranoidea | Genus | *Cornufer vitianus* CM 1498 | 12220 | 9271 | 332 | 78 | 12371 | 848.8 | 793 |
| Ranoidea | Genus | *Cornufer weberi* SLT 269 | 13207 | 10678 | 371 | 139 | 12594 | 1054.3 | 1020 |
| Ranoidea | Genus | *Cornufer wuenscheorum* RG 7751 | 11456 | 9364 | 267 | 108 | 12181 | 799.4 | 746 |
| Reduced | Reduced | *Occidozyga sp* CDS 2282 | 13450 | 2718 | 70 | 255 | 8032 | 961.3 | 817.5 |
| Reduced | Reduced | *Occidozyga sp* CDS1257 | 13736 | 2787 | 106 | 376 | 8762 | 1045.2 | 861 |
| Reduced | Reduced | *Occidozyga sp* CDS1808 | 16694 | 2853 | 111 | 337 | 12094 | 1035.8 | 844 |
| Reduced | Reduced | *Occidozyga sp* CDS2026 | 17525 | 2875 | 110 | 188 | 20351 | 1055.6 | 864 |
| Reduced | Reduced | *Occidozyga sp* CDS2045 | 19231 | 2870 | 116 | 152 | 12048 | 1044.3 | 857 |
| Reduced | Reduced | *Occidozyga sp* CDS270 | 19332 | 2832 | 123 | 379 | 11986 | 1080.9 | 890.5 |
| Reduced | Reduced | *Occidozyga sp* CDS3582 | 15784 | 2807 | 88 | 186 | 12054 | 1036.7 | 829 |
| Reduced | Reduced | *Occidozyga sp* CDS487 | 20810 | 2953 | 122 | 446 | 8110 | 1050.5 | 884 |
| Reduced | Reduced | *Occidozyga sp* CDS543 | 21314 | 2938 | 134 | 251 | 7344 | 998.1 | 861 |
| Reduced | Reduced | *Occidozyga sp* RMB11020 | 16437 | 2862 | 74 | 287 | 8238 | 1068.1 | 908 |
| Reduced | Reduced | *Occidozyga sp* RMB15467 | 14586 | 2847 | 94 | 227 | 11997 | 991.8 | 820 |

| | | | | | | | | | |
|---|---|---|---|---|---|---|---|---|---|
| Reduced | Reduced | *Occidozyga sp* RMB16321 | 23856 | 2935 | 123 | 336 | 12276 | 1103.0 | 908 |
| Reduced | Reduced | *Occidozyga sp* RMB16436 | 20593 | 2891 | 118 | 253 | 12134 | 1080.2 | 883 |
| Reduced | Reduced | *Occidozyga sp* RMB17235 | 25179 | 2983 | 125 | 431 | 12180 | 1140.9 | 950 |
| Reduced | Reduced | *Occidozyga sp* RMB20282 | 25553 | 2868 | 128 | 352 | 12774 | 1088.1 | 887 |
| Reduced | Reduced | *Occidozyga sp* RMB20559 | 13377 | 2769 | 91 | 359 | 12118 | 1038.5 | 861 |
| Reduced | Reduced | *Occidozyga sp* RMB2133 | 11852 | 2747 | 84 | 351 | 12085 | 1057.8 | 876 |
| Reduced | Reduced | *Occidozyga sp* RMB2542 | 7766 | 2299 | 50 | 154 | 11768 | 945.4 | 735 |
| Reduced | Reduced | *Occidozyga sp* RMB3068 | 20096 | 2942 | 132 | 297 | 12079 | 1095.4 | 910 |
| Reduced | Reduced | *Occidozyga sp* RMB3127 | 17714 | 2829 | 99 | 215 | 12132 | 1056.0 | 874 |
| Reduced | Reduced | *Occidozyga sp* RMB4318 | 18594 | 2904 | 118 | 242 | 13443 | 984.3 | 847 |
| Reduced | Reduced | *Occidozyga sp* RMB4468 | 25764 | 2933 | 115 | 309 | 12153 | 1098.9 | 906 |
| Reduced | Reduced | *Occidozyga sp* RMB5396 | 19282 | 2865 | 108 | 326 | 12250 | 1058.8 | 874 |
| Reduced | Reduced | *Occidozyga sp* RMB6117 | 18603 | 2910 | 96 | 247 | 8569 | 1025.1 | 865 |
| Reduced | Reduced | *Occidozyga sp* RMB6411 | 13723 | 2804 | 101 | 434 | 12016 | 1055.3 | 868 |
| Reduced | Reduced | *Occidozyga sp* RMB7686 | 11193 | 2826 | 91 | 272 | 7881 | 1008.0 | 871 |
| Reduced | Reduced | *Occidozyga sp* RMB819 | 12873 | 2790 | 90 | 319 | 12105 | 1080.2 | 887 |
| Reduced | Reduced | *Occidozyga sp* RMB8750 | 21191 | 2894 | 119 | 226 | 12641 | 1067.0 | 874 |
| Reduced | Reduced | *Occidozyga sp* RMB8900 | 15194 | 2910 | 103 | 403 | 12119 | 1095.2 | 908 |
| Reduced | Reduced | *Occidozyga sp* RMB9600 | 25680 | 2974 | 119 | 172 | 12100 | 1097.2 | 910 |
| Hyloidea | Superfamily | *Allobates femoralis* WED 55592 | 24894 | 5662 | 372 | 163 | 9924 | 852.3 | 781.5 |
| Hyloidea | Superfamily | *Anaryxus woodhousii* RMB 19577 | 22188 | 7699 | 1750 | 73 | 10914 | 1019.8 | 962 |
| Hyloidea | Superfamily | *Ansonia sp* RMB 10371 | 24794 | 7810 | 2718 | 180 | 8799 | 1103.4 | 1055 |
| Hyloidea | Superfamily | *Barycholos pulcher* LAC 692 | 16724 | 5907 | 451 | 122 | 11654 | 1014.0 | 954 |
| Hyloidea | Superfamily | *Bolitaglossa palmata* LAC 1154 | 13790 | 535 | 20 | 88 | 3642 | 762.6 | 720 |
| Hyloidea | Superfamily | *Bufo valliceps* KU 203846 | 43113 | 8232 | 3475 | 100 | 8605 | 1244.2 | 1178 |
| Hyloidea | Superfamily | *Ceratophrys cornuta* WED 57689 | 20379 | 7889 | 1073 | 84 | 10112 | 1054.4 | 994 |
| Hyloidea | Superfamily | *Craugastor rupinius* JES 2401 | 19238 | 6570 | 1288 | 167 | 8343 | 1135.3 | 1058 |
| Hyloidea | Superfamily | *Dendrobates histrionicus* LAC 2539 | 11997 | 5459 | 343 | 120 | 8204 | 851.2 | 788 |
| Hyloidea | Superfamily | *Eluetherodactylus sp* GLOR 6202 | 17099 | 6733 | 963 | 139 | 11649 | 1163.2 | 1102 |

| | | | | | | | | | |
|---|---|---|---|---|---|---|---|---|---|
| Hyloidea | Superfamily | *Gastrotheca marsupiata* WED 58521 | 13538 | 6559 | 797 | 108 | 7923 | 865.5 | 800 |
| Hyloidea | Superfamily | *Hyalinobatrachium fleischmanni* MZUTI 3621 | 17393 | 6705 | 1316 | 201 | 8343 | 1059.0 | 989 |
| Hyloidea | Superfamily | *Hyloscirtus phyllognathus* WED 58378 | 16567 | 7157 | 809 | 98 | 13871 | 1058.8 | 1006 |
| Hyloidea | Superfamily | *Leptobrachium hainensis* KUFS 194 | 8193 | 2418 | 75 | 104 | 6717 | 828.8 | 785 |
| Hyloidea | Superfamily | *Leptodactylus pentadactylus* WED 55494 | 22665 | 7677 | 864 | 79 | 16967 | 1078.1 | 1029 |
| Hyloidea | Superfamily | *Nyctimystes infrafrenatus* SLT 771 | 25534 | 7940 | 978 | 83 | 11291 | 1105.6 | 1047 |
| Hyloidea | Superfamily | *Oreobates quizensis* WED 59885 | 12446 | 5870 | 953 | 112 | 8416 | 913.2 | 842 |
| Hyloidea | Superfamily | *Phyrnohyas venulosa* WED 55450 | 20170 | 7320 | 859 | 157 | 9389 | 1081.0 | 1023 |
| Hyloidea | Superfamily | *Physalamus pustulatus* LAC 1642 | 25150 | 7259 | 969 | 102 | 7745 | 1033.5 | 982 |
| Hyloidea | Superfamily | *Pristimantis w-nigrum* WED 53045 | 26099 | 7576 | 1521 | 107 | 11463 | 1101.7 | 1040 |
| Hyloidea | Superfamily | *Spea bombifrons* RMB 19586 | 21131 | 4328 | 246 | 88 | 7441 | 1067.5 | 1016 |
| Hyloidea | Superfamily | *Strabomantis sulcatus* LAC 841 | 17898 | 6754 | 1459 | 121 | 8466 | 1002.3 | 931 |
| Hyloidea | Superfamily | *Telmatobius truebae* WED 56933 | 28122 | 8337 | 1916 | 195 | 8743 | 1282.1 | 1226 |
| Hyloidea | Superfamily | *Vitreorana castroviejoi* JMG 33 | 22072 | 7183 | 2434 | 78 | 8044 | 1059.6 | 999 |

**Figure S1.** A summary of the data obtained for each sample, shown at each phylogenetic scale. The total number of markers in (A) are those obtained after matching to the target loci and removal of paralogs. In (B), the proportion is the sample total number of markers divided by the number of target markers. Finally, in (C) the total number of megabase pairs are the number of aligned bases before (light green) and after trimming (dark green).

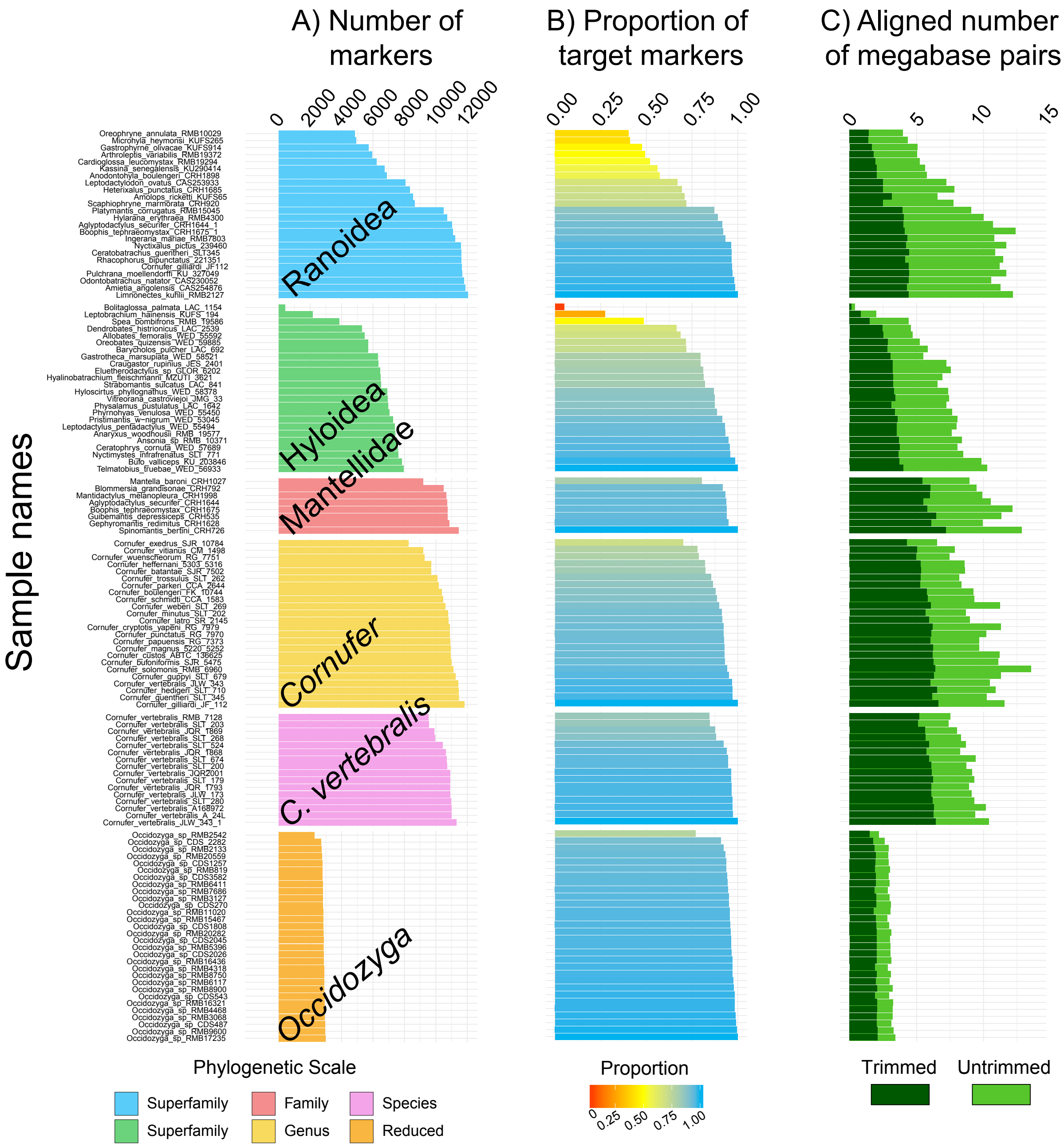
